# PRDM16 co-operates with LHX2 to shape the human brain

**DOI:** 10.1101/2023.08.12.553065

**Authors:** Varun Suresh, Bidisha Bhattacharya, Rami Yair Tshuva, Miri Danan Gotthold, Tsviya Olender, Mahima Bose, Saurabh J. Pradhan, Bruria Ben Zeev, Richard Scott Smith, Shubha Tole, Sanjeev Galande, Corey Harwell, José-Manuel Baizabal, Orly Reiner

**Affiliations:** Departments of Molecular Genetics and Molecular Neuroscience, Weizmann Institute of Science, Rehovot, Israel; Department of Biological Sciences Tata Institute of Fundamental Research, Mumbai, India; Edmomd and Lilly Safra Pediatric Hospital, Sheba Medical Center and Tel Aviv School of Medicine, Tel Aviv University, Ramat Aviv 69978, Israel; Department of Pharmacology at Northwestern University, Feinberg School of Medicine, Chicago, IL, USA; Chromatin Biology and Epigenetics Laboratory, Biology department, Indian Institute of Science Education and Research Pune, India; Center of Excellence in Epigenetics, Department of Life Sciences, Shiv Nadar University, India; Eli and Edythe Broad Center of Regeneration Medicine and Stem Cell Research, University of California, San Francisco, San Francisco, CA, US; Weill Institute for Neuroscience, San Francisco, CA, USA; Department of Neurology, University of California, San Francisco, San Francisco, CA, USA; Department of Biology, Indiana University, Bloomington, IN, USA; Institute of Molecular Biotechnology of the Austrian Academy of Sciences, Vienna Biocenter, Vienna, Austria

## Abstract

PRDM16 is a dynamic transcriptional regulator of various stem cell niches, including adipocytic, hematopoietic, cardiac progenitors, and neural stem cells. PRDM16 has been suggested to contribute to 1p36 deletion syndrome, one of the most prevalent subtelomeric microdeletion syndromes. We report a patient with a de novo nonsense mutation in the PRDM16 coding sequence, accompanied by lissencephaly and microcephaly features. Human stem cells were genetically modified to mimic this mutation, generating cortical organoids that exhibited altered cell cycle dynamics. RNA sequencing of cortical organoids at day 32 unveiled changes in cell adhesion and WNT-signaling pathways. ChIP-seq of PRDM16 identified binding sites in postmortem human fetal cortex, indicating the conservation of PRDM16 binding to developmental genes in mice and humans, potentially at enhancer sites. A shared motif between PRDM16 and LHX2 was identified and further examined through comparison with LHX2 ChIP-seq data from mice. These results suggested a collaborative partnership between PRDM16 and LHX2 in regulating a common set of genes and pathways in cortical radial glia cells, possibly via their synergistic involvement in cortical development.

## Introduction

PRDM16 (PR/SET Domain 16), a versatile protein, influences transcription by forming complexes with various transcription factors and regulators and through its histone methyl transferase activities [1–5]. This dynamic regulator has emerged as a key player in several stem cell niches, including brown fat precursors, hematopoietic cells, cardiac progenitors, palate and craniofacial precursors, and embryonic and adult neural stem cells [1,6–18]. Extensive research has demonstrated the significance of PRDM16 in various biological processes. Mutations in PRDM16 have been identified in patients with cardiomyopathy, and zebrafish models have successfully recapitulated similar phenotypes [19,20]. Notably, chromosomal rearrangements involving PRDM16 contribute to hematological malignancies, such as myelodysplastic syndromes (MDS) and acute myeloid leukemia (AML). At the same time, aberrant expression of the short variant of PRDM16 (PRDM16/MEL1S) has been implicated in a range of hematological malignancies, including T-cell leukemia [21–23].

*PRDM16* is located in the telomeric region of chromosome 1 (1p36), which is susceptible to *de novo* deletions [24–30], while duplications and triplications in this locus are less frequent [28,31–33]. The 1p36 deletion syndrome, characterized by variable deletion sizes, is considered one of the most prevalent subtelomeric microdeletion syndromes, affecting approximately 1 in 5000 live births [24,34]. Phenotypic severity does not correlate strongly with deletion size, suggesting complex interactions within the haploinsufficient protein products [30]. Common manifestations of 1p36 deletion syndrome include hypotonia, severe developmental delay, growth retardation, microcephaly, obesity, craniofacial dysmorphism, cardiac malformations, cardiomyopathy, and seizures [35–37]. Among the genes in this region, *SKI, GABRD, RERE, PRDM16*, and *KCNAB2* have been proposed as contributors to the observed syndromic phenotypes [38].

In the developing mouse brain, PRDM16 exhibits robust expression in radial glia (RG) progenitors, serving as one of the conserved core RG genes shared between humans and mice [11,14,17,39–43]. Additionally, PRDM16 is expressed in adult neural stem cells [15]. Studies using *Prdm16* null and Nestin-regulated postnatal deletion mice have revealed a reduced number of phospho-histone-positive cells and a diminished capacity to form neurospheres [11,15]. Postmitotic radial glia progeny during brain development resides in the ventricular zone before transitioning into multipolar cells that migrate towards the intermediate zone [43]. PRDM16 expression gradually declines as progenitors progress through differentiation stages from the ventricular zone to the intermediate zone [14,41]. Knockdown and overexpression experiments have demonstrated the role of PRDM16 in regulating the development of postmitotic multipolar cells [14]. Knockdown of *Prdm16* induces the expression of markers associated with the terminal multipolar stage, while overexpression expands the population of early multipolar cells (NeuroD1+) [14]. Furthermore, PRDM16 and NeuroD1 are crucial for regulating PGC1a activity, essential for controlling global oxidative metabolism, which undergoes significant changes during the multipolar transition [44].

Much is known regarding the functions of PRDM16 in the developing mouse brain, yet its roles in the developing human brain still need to be explored. Our study was motivated by detecting a patient with a *de novo* nonsense mutation in *PRDM16* exhibiting lissencephaly and microcephaly features. We introduced the mutation in human stem cells in both homozygous and heterozygous manner and generated human cortical organoids. PRDM16 levels affected the cell cycle, and RNA-seq indicated changes in cell adhesion and WNT-signaling, which were confirmed by immunostaining. We identified PRDM16 binding sites across the genome of radial glia cells in postmortem human fetal samples. Our results indicate high conservation of PRDM16 binding to mouse and human developmental genes, possibly at enhancers [39,41]. The top detected motif matched LHX2, so we compared our data with recently acquired LHX2 ChIP-seq data from the mouse. Our studies show that PRDM16 and LHX2 collaborate in regulating a common set of genes and pathways in cortical RG, indicating possible synergistic action by these proteins.

## Results

*PRDM16* is expressed in apical and basal progenitors in the developing brain of humans **(Fig. 1A-B)** and mice [39,41]. To better understand the expression of *PRDM16* RNA, we used publicly available single-cell RNAseq data from early human embryonic brains at 5 – 14 post-conceptional weeks (PCW) [45]. PRDM16 was one of the genes that showed higher expression in late radial glia from the developing pallium (EMX1+).

**Fig. 1.**
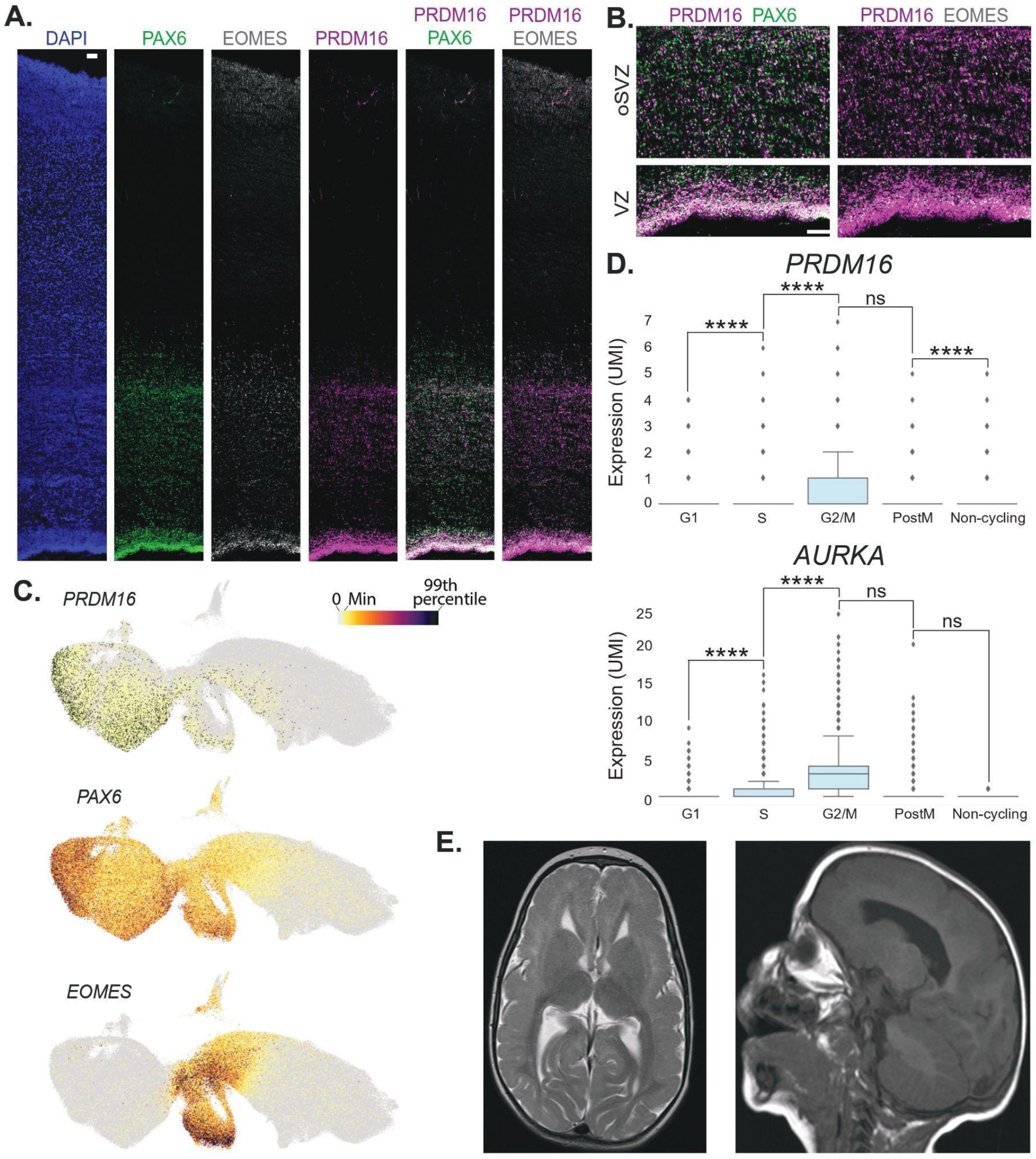
PRDM16 expression and patient data A-B. Immunostaining of human embryonic brain sections of the cingulate cortex (GW20-22). **Left to right in A.** DAPI, PAX6, EOMES, PRDM16, and merged images of PRDM16 staining with PAX6 and EOMES. **B.** Zoom-in images of the double immunostainings in the ventricular zone (VZ) and the outer subventricular zone (oSVZ). The scale bars in A and B are 100 µm. **C.** UMAP of pallial excitatory neuron lineage at 5-14 PCW from [45] colored by selected gene expression (the color scale on the top right ranges from gray (0) to black (99 percentile). **D.** Box plots showing the gene expression *PRDM16* (top) and *AURKA* (bottom) for each progenitor state in radial glia at 5-14 p.c.w, the Y axis indicates expression levels in UMI (*EMX1*+ clusters, data from [45]) (Two-sided Mann-Whitney-Wilcoxon test with Bonferroni correction is used to test the significance. ns: not significant; P-value: **** <=0.0001). **E.** Two MRI images from the patient exhibiting bilateral frontal pachygyria and microcephaly, T2 transverse (left) and sagittal (right).

*PRDM16* was expressed in both neuronal intermediate progenitor cells (nIPCs) and RG (expressing EOMES and HES1, respectively) **(Fig. 1C)** in agreement with previously reported data [41]. Analyzing these radial glial cells from 5-14 PCW revealed that *PRDM16* is expressed in a cell-cycle-dependent manner somewhat similar to Aurora Kinase (*AURKA)* **(Fig. 1D)**. There is a significant increase in its expression levels when the cells undergo G1→ S transition and a decrease following mitosis (post-M stage).

PRDM16 has a possible role in the 1p36 deletion syndrome, and we identified a single microcephaly/lissencephaly patient. The identified patient exhibited severe developmental delay, intellectual disability, intractable epilepsy, and hypotonia. MRI detected bilateral frontal pachygyria and microcephaly **(Fig. 1E and Supplementary** Fig. 1A). Seizures consistent with prolonged staring followed by head drops were noticed at two years of age. Electroencephalogram (EEG) showed diffuse generalized and multifocal spike waves aggravated in sleep (consistent with electrical status in sleep). From the onset of seizures till today, the child has had frequent seizures, including atypical absence and short generalized tonic-clonic seizures resistant to a long list of anticonvulsants and the ketogenic diet. We applied exome sequencing to the child and his two parents to identify a possible genetic mutation underlying the disease. We identified a single candidate mutation in the child, a *de novo* frame-shift deletion in the *PRDM16* gene (a deletion of T at chr1: 3396566 (hg38), Leu217fs, NM_199454). No other candidate mutations were identified from the child exome data. The sequence was verified by Sanger sequencing **(Supplementary** Fig. 1B **up)**.

We employed a brain organoid-based strategy to investigate how this mutation affects the development of the human forebrain. We inserted the patient-specific mutation in human embryonic stem cells (hESCs). We used a double-nickase approach using a single-strand oligonucleotide repair to reduce possible off-targets. Homozygous and heterozygous lines were generated and validated by sequencing **(Supplementary** Fig. 1B **down)**. We generated cortical organoids from one homozygous and one heterozygous line based on Sasai’s protocol [46,47]. On day 32 of the organoid culture, the organoids expressed progenitor markers such as PAX6, SOX2, and oSVZ markers such as HOPX **(Fig. 2A)**. We noted a similar percentage of SOX2+ progenitors from the total number of cells in WT and mutant organoids **(Supplementary** Fig. 1C-D). In contrast, the heterozygous and the homozygous *PRDM16* mutant organoids exhibited a significant increase of PAX6-positive progenitors **(Fig. 2B left)**.

**Fig. 2.**
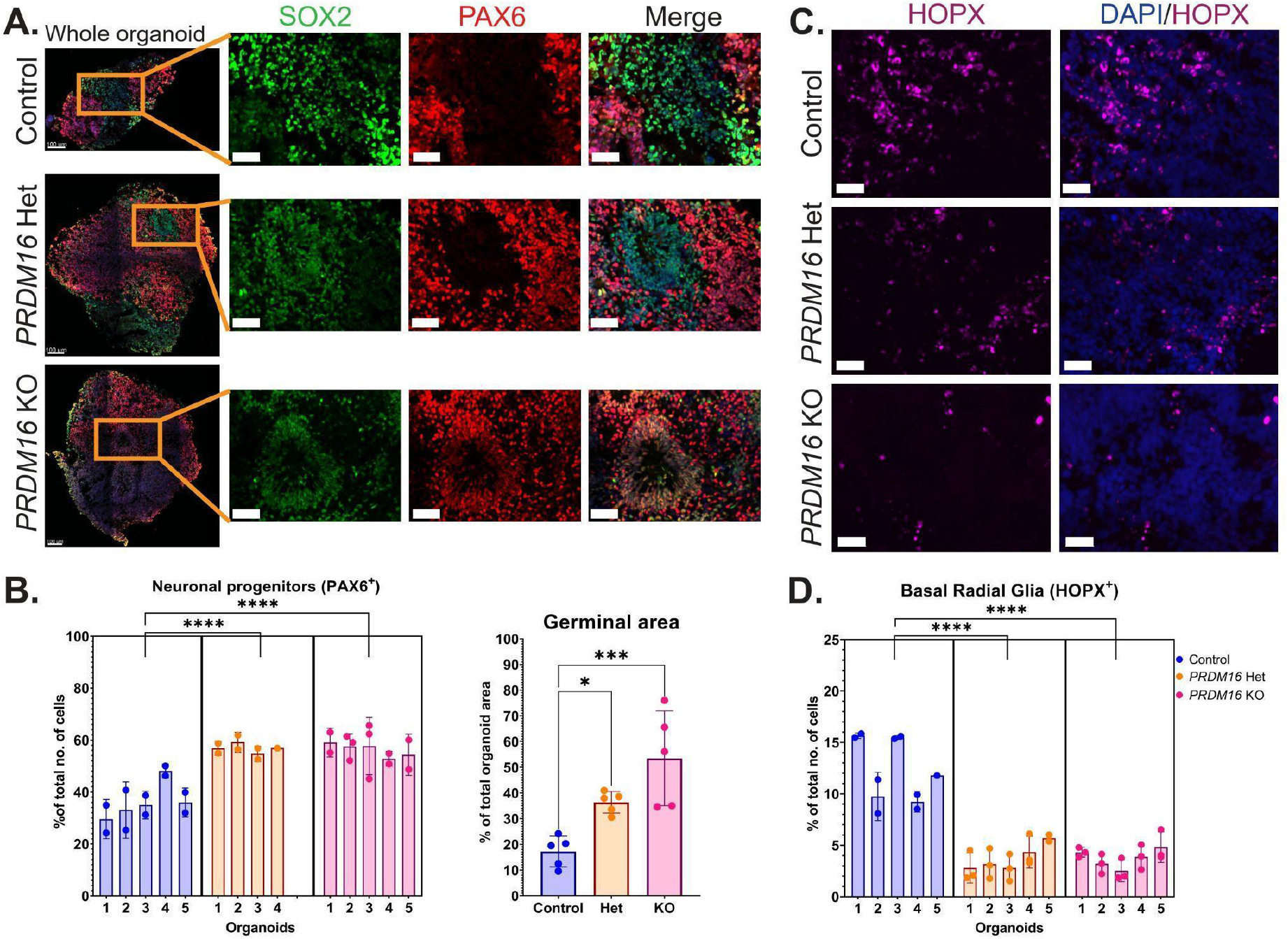
Cortical progenitors in *PRDM16* mutant human brain organoids A. Immunohistochemistry on 12 µm cryosections of day 32 control and *PRDM16* mutant cortical organoids showing the cortical progenitor markers – SOX2 and PAX6 (imaged using 25X objective lens, scale bar represents 50 µm in insets) **B** (left). Percentage of total number of cells that are SOX2^+^ and PAX6^+^ neuronal progenitors respectively, in day 32 control and PRDM16 mutant cortical organoids (Nested One-Way ANOVA, n = 5, α = 0.05, ns: not significant; p-values: * <= 0.05; ** <= 0.01; *** <=0.001; **** <=0.0001, x-axis represents the number of organoids quantified, each data point on the graph represents an image of the organoid along its z-axis) and **B** (right) the total area of proliferative germinal zones are represented as a percentage of total organoid area in control and *PRDM16* mutant organoids (Ordinary One-Way ANOVA, n = 5, α = 0.05) **C.** Immunohistochemistry on 12 µm cryosections of day 32 control and *PRDM16* mutant cortical organoids showing the basal radial glia markers – HOPX (imaged using 25X objective lens, scale bar represents 30 µm) **D.** Percentage of total number of cells that are HOPX^+^ in day 32 control and PRDM16 mutant cortical organoids (Nested One-Way ANOVA, n = 5, α = 0.05)

Furthermore, we noted a change in the relative appearance of germinal zones in the organoids from the different genotypes. Our measurements revealed an increase in the germinal zone area in mutant organoids compared with the control organoids, with a significant increase in the heterozygous (p<0.05) and a further considerable elevation in the homozygous mutant (p<0.01) **(Fig. 2B right)**. However, the percentage of the HOPX+ progenitors in both *PRDM16* mutants was significantly reduced **(Fig. 2C-D)**.\

Compared with the control, analysis of proliferation and cell cycle markers revealed an increase in KI67+ proliferating cells in the *PRDM16* KO organoids. While no change in M-phase was observed (pHH3+ cells), we noted more cells in the S-phase in the KO organoids (EdU positive). In these KO organoids, more cells were exiting the cell cycle (KI67-/EdU+) (**Fig. 3)**. Overall, these results suggest that the loss of *PRDM16* promotes an increase in the self-renewal of PAX6-positive RG while concomitantly increasing the rate of differentiation (higher cell cycle exit) of the RG-progeny.

**Fig. 3.**
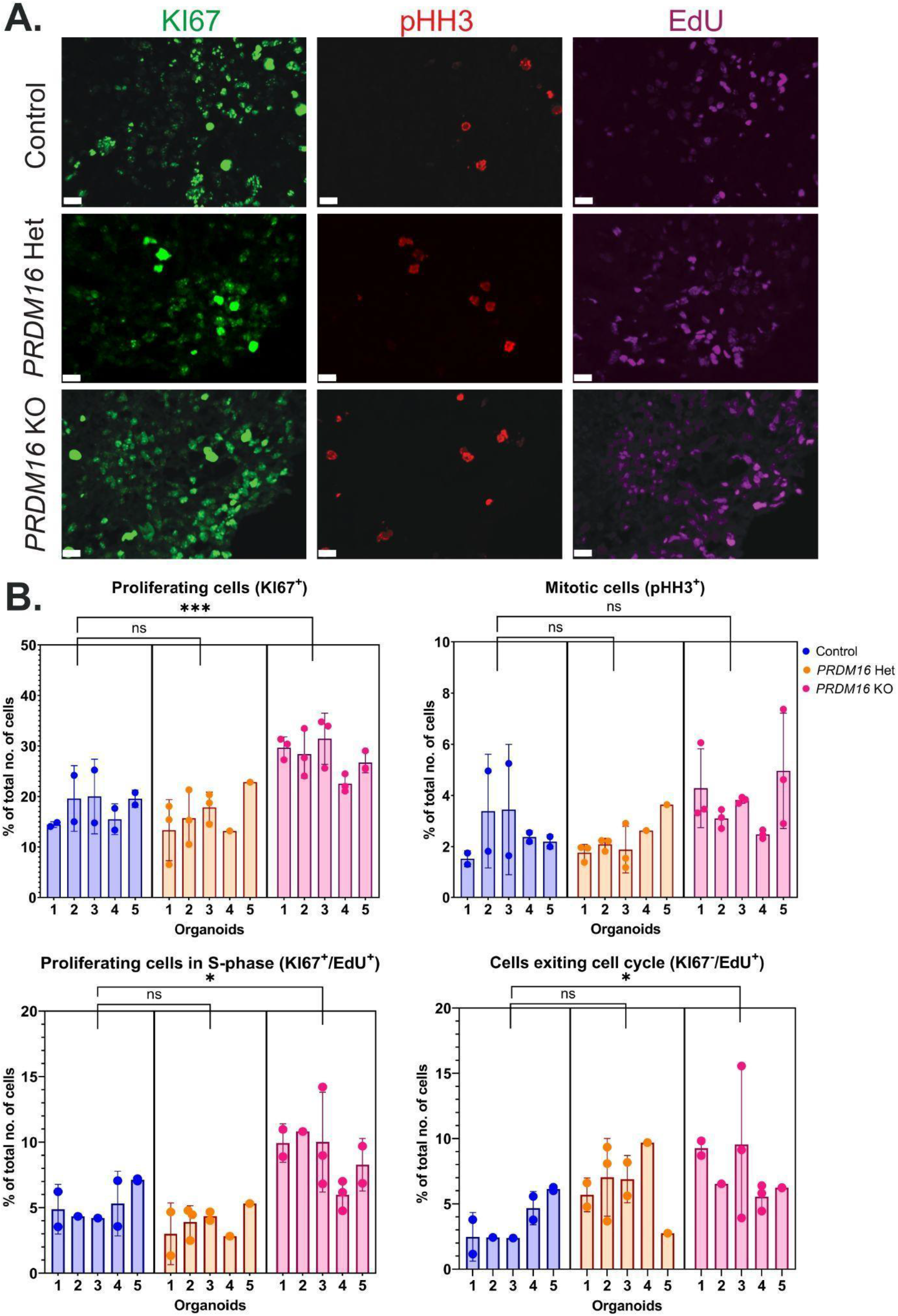
Proliferating cells in *PRDM16* mutant human brain organoids A. Immunohistochemistry on 12 µm cryosections of day 32 control and PRDM16 mutant cortical organoids showing the proliferative markers – KI67, pHH3, and EdU (imaged using 25X objective lens, scale bar represents 15 µm) **B.** Percentage of the total number of cells that are proliferative (KI67^+^), mitotically-active (pHH3^+^), proliferative cells undergoing S-phase (KI67^+^/EdU^+^) and cells that are exiting the cell cycle (KI67^-^/EdU^+^) respectively, in day 32 control and PRDM16 mutant cortical organoids (Nested One-Way ANOVA, n = 5, α = 0.05)

To further improve our understanding of the effects of the loss of function of PRDM16, we characterized the transcriptome. We conducted RNAseq on the homozygous mutant cortical organoids and the corresponding control on day 32. RNA-seq analysis resulted in the identification of 3203 differentially expressed genes (DEGs). Of these, 2205 down-regulated genes and 998 genes were up-regulated in the mutant versus control organoids (Fold change > 2 and P-adj < 0.05) **(Fig. 4A, Supplementary** Fig. 2B-D**, Supplementary Table S1)**. The Gene Ontology Biological processes (GO: BPs) and Kyoto Encyclopedia for genes and genomes (KEGG) pathway analysis of the DEGs revealed multiple developmental pathways affected, such as pathways related to WNT signaling, cell adhesion, and ECM-receptor interactions **(Fig. 4B and Supplementary** Fig. 3A). Of particular interest were genes belonging to the WNT pathway, a prominent developmental pathway that controls progenitor proliferation, cell cycle exit, and neuronal morphology in the developing mammalian brain, which is possibly implicated in neurodevelopmental disorders [48,49]. We detected 33 DEGs belonging to the WNT pathway genes. The downregulated WNT pathway genes included receptors such as the *FZD* family of genes, *ROR2, LGR6*, and ligands such as *RSPO4, WNT8A*, and *WNT11*. We further noted the downregulation of WNT antagonists, such as *DKK1&2* and *WIF1*, was coupled with the upregulation of multiple WNT ligands, such as *WNT 2B, 4, 7B*, and *8B*. In addition, we noted elevated expression of receptors such as *FZD2* and *LGR5* and agonists such as *WLS* and *DKK4* **(Fig. 4C)**. Consistent with these observations, the intensity of nuclear CTNNB1, the canonical WNT effector molecule, was increased in *PRDM16* mutant organoids **(Fig. 4E-F)**. Accordingly, we observed an overall increase in the percentage of LEF1+ cells in the mutant organoids, suggesting WNT signaling activation.

**Fig. 4.**
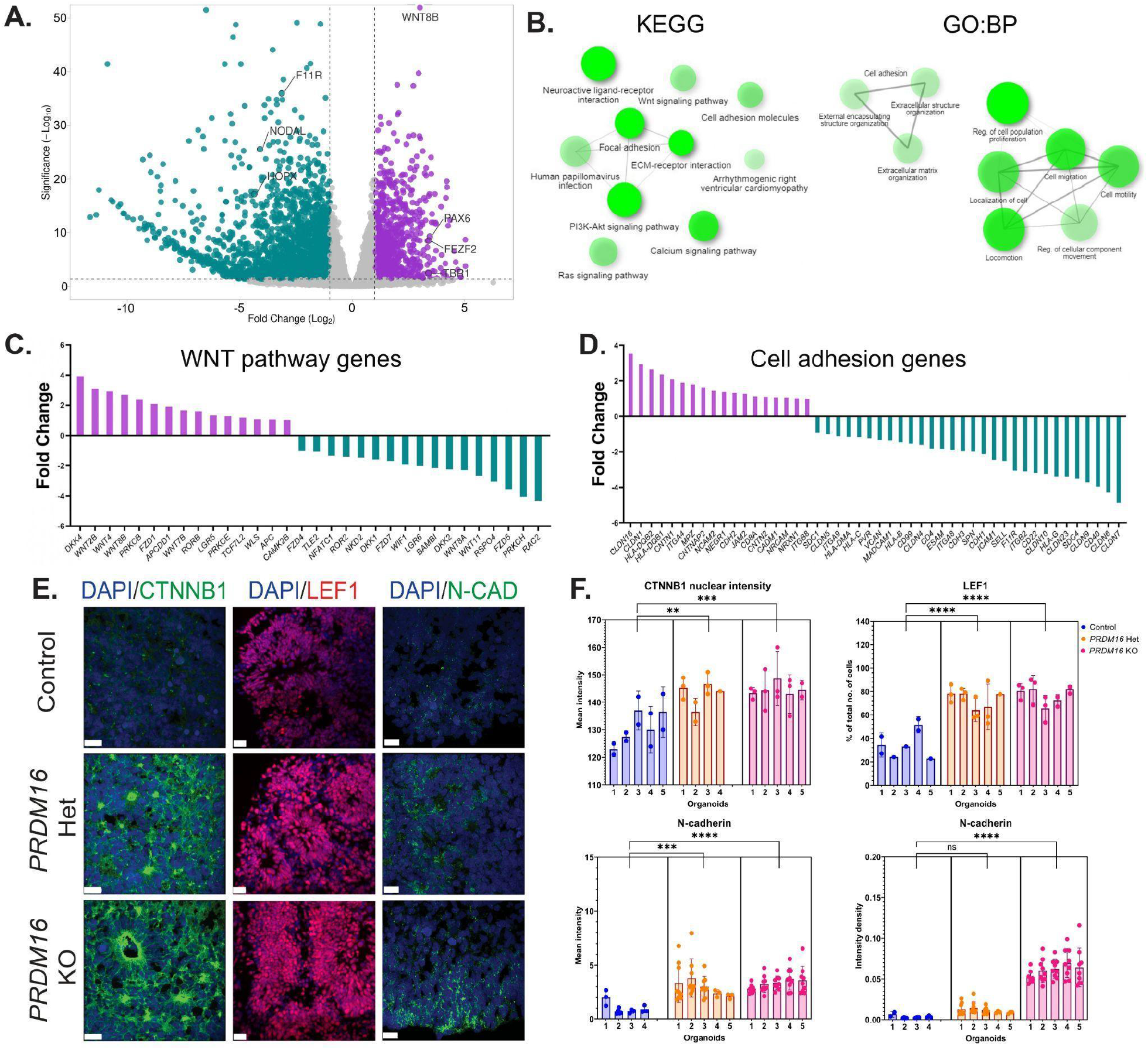
Transcriptomic analysis of PRDM16 mutant organoids identifies dysregulated WNT signaling and cell adhesion. **A.** Volcano plot shows DEGs identified in *PRDM16* mutant organoids vs control—B. Gene ontology: Biological processes and KEGG pathway analysis of DEGs identified in A, the data was plotted using the “Network” plot of ShinyGO [50]. **C-D.** Fold changes of genes belonging to WNT signaling and cell adhesion biological processes depict multiple dysregulated genes. **E.** Immunohistochemistry on 12 µm cryosections of day 32 control and PRDM16 mutant cortical organoids showing two markers of Wnt signaling; CTNNB1 and LEF1 (imaged using 25X objective lens, scale bar represents 20 µm) and N-cadherin cell adhesion molecule (imaged using 63X objective lens, scale bar represents 20 µm) **F.** Mean intensity of CTNNB1 inside the nuclei of cells in control and *PRDM16* mutant cortical organoids (Nested One-Way ANOVA, n = 5, α = 0.05), percentage of total number of cells that are LEF1^+^, on day 32 control and PRDM16 mutant cortical organoids (nested one-way ANOVA, n = 5, α = 0.05), mean intensity and intensity density of N-cadherin at the boundary of the cells in control and PRDM16 mutant cortical organoids (nested one-way ANOVA, ncontrol = 4, nPRDM16 KO = 5, α = 0.05).

Pathway analysis also suggested an alteration in the expression of genes associated with cell adhesion and extracellular matrix. To support these data, we immunostained the organoids using anti-N-CADHERIN antibodies and observed an overall increase in the mutant organoids’ mean intensity and area density compared to the control ones **(Fig. 4E-F)**. These results indicate that multiple components of the WNT pathway and cell adhesion processes are dysregulated in the mutant organoids.

To investigate the potential gene regulatory networks controlled by PRDM16, we performed ChIP-seq (Chromatin Immunoprecipitation and sequencing) from dissected cortical progenitor regions of human fetal samples at gestational week (GW) 15 and 22. Our analysis of PRDM16 occupancy profiles identified 7155 high-confidence peaks representing highly reproducible PRDM16 binding sites in the two human samples **(Fig. 5A and Supplementary Table S2)**. By comparing the human PRDM16 ChIP-seq to previously published PRDM16 ChIP-seq data in the mouse cortex ([39,41], see methods), we revealed that 986 peaks of the regions identified in the human, ChIPseq showed an overlapping peak in the mouse PRDM16 occupancy data **(Fig. 5B, C)**. Moreover, most overlapping PRDM16 peaks between human and mouse were bound to either intronic or intergenic regions, suggesting that PRDM16 also binds to developmental enhancers in the human brain [40]. To identify the possible direct targets of PRDM16, we compared the ChIP-seq generated from human tissues with the DEGs identified in the brain organoids upon loss of PRDM16, revealing a set of 622 genes **(Supplementary** Fig. 4A**, Supplementary Table S3)**. The GO: BPs terms associated with the genes directly regulated by PRDM16 were related to neurogenesis, cell fate commitment, and axon development, suggesting a role for PRDM16 in regulating these processes in the human cortex **(Fig. 5D)**. Among the nearest genes to PRDM16 binding sites that are also differentially expressed in mutant organoid, we found *FEZF2, ROBO2, RBFOX1,* and *SOX5* **(Fig. 5E)**. All of these genes have been shown to play critical roles during neurogenesis in the mouse, and some of them, such as *FEZF2,* were shown to be PRDM16 targets during mouse development [39,41] **(Supplementary Table S1, S3)**. We investigated if the loss of PRDM16 also results in increased neurogenesis in our organoids by staining for the pan-neuronal marker MAP2 and layer six neuronal marker TBR1. We observed that the number of TBR1+ cells in the mutant organoids increased. The number of MAP2+ cells is increased only in the homozygous mutant organoids. However, the number of CTIP2+ cells did not vary **(Fig. 5F-G, Supplementary** Fig. 4B**-C)**. These results suggest PRDM16 may affect neuronal fate acquisition during human brain development.

**Fig. 5.**
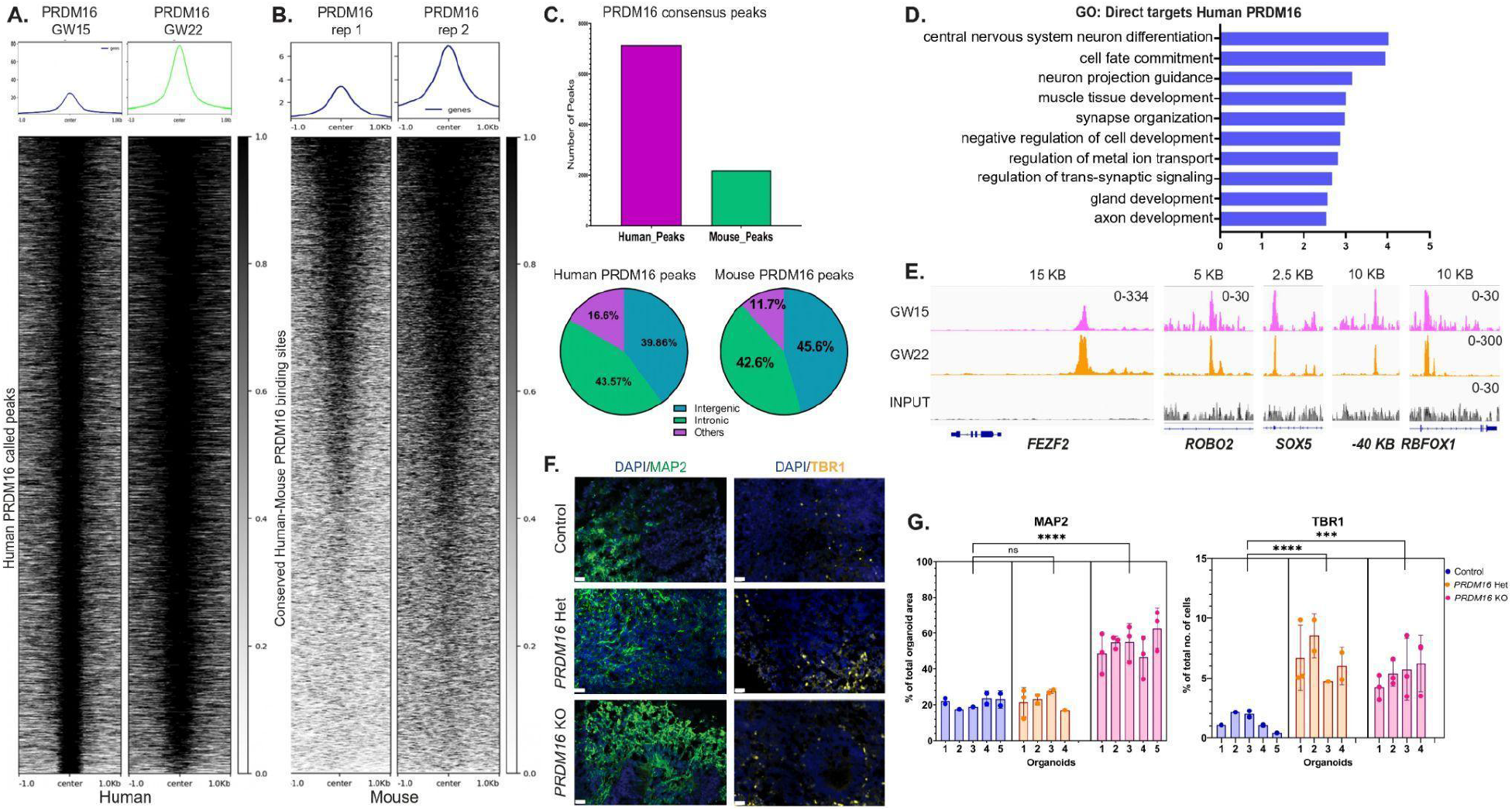
A. PRDM16 occupies and regulates multiple genes that regulate cortical neurogenesis. Heatmap and profile plots of the Human PRDM16 ChIP-seq data show PRDM16 consensus binding sites. **B.** Mouse PRDM16 ChIP-seq data (obtained from [41]) plotted on regions identified in A after a human-to-mouse LiftOver conversion. **C.** Annotations of Mouse and Human PRDM16 ChIP-seq data reveal the number of PRDM16-occupied regions in both species. **D.** Gene Ontology Biological Processes directly regulated by PRDM16 identified by genes occupied by PRDM16 and dysregulated in the organoid. **E.** Select IGV tracks of neuronal genes occupied by PRDM16 in the human fetus, namely - *FEZF2* (chr3:62. 327, 944-62, 384, 680), *ROBO2* (chr3:77, 228, 242-77, 341, 714), *SOX5* (chr12:24, 267, 871-24, 574, 676) and *RBFOX1* (chr16:7, 669, 904-7, 877, 008). **F.** Immunohistochemistry on 12 µm cryosections of day 32 control and *PRDM16* mutant cortical organoids showing immunostainings for MAP2 and TBR1 (imaged using a 25X objective, scale bar represents 20 µm) **G.** Percentage of the total organoid area that has processes of MAP2+ neurons in control and *PRDM16* mutant organoids (nested one-way ANOVA, n = 5, α = 0.05), percentage of the total number of cells that are TBR1+ neurons in day 32 control and *PRDM16* mutant cortical organoids (nested one-way ANOVA, n = 5, α = 0.05)

Motif analysis of human PRDM16 binding sites revealed that the consensus DNA binding motif for LHX2 was the most enriched one. 34.88% of all peaks identified in our ChIP-seq analysis contained the LHX2 motif. In comparison, only 5.17% of peaks contained PRDM9 motifs, and 2.94% of peaks had the PRDM14 motifs, and the PRDM16 motif was not detected as statistically significant in this dataset. Furthermore, we confirmed that LHX2 was the top enriched motif within PRDM16 binding sites in the mouse embryonic cortex, consistent with previous reports [39,41], where over 40% of all consensus peaks across mice and human regions consisted of LHX2 motifs. **(Fig. 6A)**. The transcription factor LHX2 is well-known for regulating multiple aspects of cortical development in the mouse [51]. Loss of LHX2 in the mouse leads to precocious differentiation of progenitors into neurons, with a remarkable increase in FEZF2 in the cortical plate [52]. To investigate the molecular mechanisms by which LHX2 controls neurogenesis and cell fate specification, we performed transcriptomic analysis of *Lhx2* cKO mouse cortex at E12.5 (embryonic day 12.5); one day after the gene was lost entirely (see methods). RNA-seq analysis revealed a set of 744 down-regulated genes and 852 up-regulated genes (FDR< 0.05 and |fold change| > 1.5) **(Fig. 6B, Supplementary Table S4)**. GO BP analysis of DEGs showed an enrichment of genes associated with neuronal differentiation, an observation that is consistent with the *Lhx2* mutant phenotype in which neurons are produced precociously [52–55] **(Fig. 6C).** Our results demonstrate a considerable overlap in the genes that both these factors regulate. We found 449 genes common to both DEG sets of mouse *Lhx2* cko and *PRDM16* mutant organoids, which is highly significant based on hypergeometric statistics showing a representation factor 55.3 with a P-value < 0.00001 (see methods) **(Fig. S4D).** These results suggest that LHX2 could be acting in concert with PRDM16, sharing a regulatory network to regulate neurogenesis in the mouse cortex. To test this hypothesis, we performed ChIP-seq for LHX2 in the E12.5 mouse cortex and identified 2222 binding sites that resulted in the identification of 1909 genes associated with these peaks (**Fig. 6D and Supplementary Table S5)**. We discovered that 447 genes were occupied by both LHX2 and PRDM16 in the mouse **(Supplementary** Fig. 4E**, Supplementary Table S7)**. Single-cell RNA expression analysis of *PRDM16* and *LHX2* in the human cortex suggested that both of these molecules are co-expressed in cortical progenitors across multiple stages of the cell cycle (**Fig. 6E**). Inspection of mouse LHX2 and PRDM16 ChIP-seq in the mouse tracks confirmed co-binding of these factors to cis-regulatory regions near neurogenic genes, such as *Fezf2, Rbfox1, and Wls* **(Fig. 6F and Supplemental Table S4)**.

**Fig. 6.**
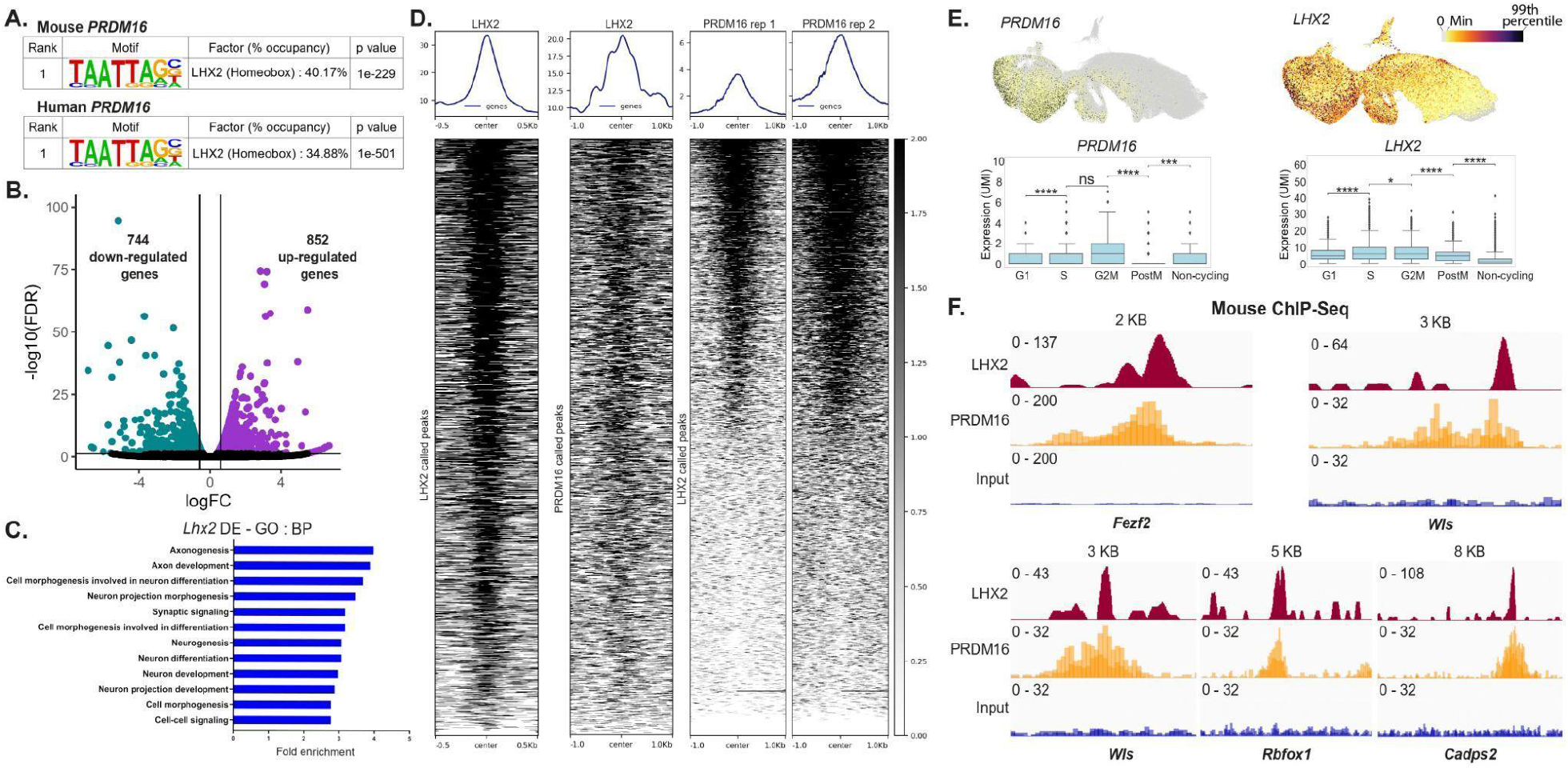
A. Motif analysis of PRDM16 mouse and human ChIP-seq data reveals the LHX2 motif as the top candidate. (a list of the top 10 candidates is available in S5)**. B.** Volcano plot of RNA-seq data shows 744 genes downregulated and 852 genes upregulated upon *Lhx2* loss in the E12.5 mouse. **C.** Gene ontology Biological Processes of the DEGs identified in B show multiple processes related to neurogenesis affected in the *Lhx2* mutant. **D.** Heat map depicts LHX2 occupancy at E12.5 in the mouse, LHX2 occupancy on PRDM16 peaks in the mouse, and heatmaps depict replicates of PRDM16 ChIP-seq data on LHX2 called peaks. **E.** UMAPs depict *PRDM16* and *LHX2* expression in the EMX1+ cells of developing human brains; bar plots indicate the expression quantified in G1, S, G2M, post-M, and cycling cells [45]. **F.** IGV tracks show common LHX2 and PRDM16 binding sites; the peaks are around 5 KB downstream of *Fezf2* (chr14:12,335,752-12, 343, 752), while all other peaks are in the intronic regions of the respective genes shown namely - *Wls* (chr3:159, 846, 672-159, 858, 135), *Rbfox1* (chr16:6, 950, 408-6, 978, 545) and *Cadps2* (chr6:23, 260, 534-23, 303, 435).

Next, we sought to identify the common gene regulatory networks controlled by PRDM16 and LHX2 in the cortex using the following stringent criteria: 1. The gene should be associated with a PRDM16 peak in the human ChIP-seq data (4054 genes); 2. The gene must also have an associated peak in the mouse PRDM16 ChIP-seq data (1354) genes, 3. The gene is dysregulated upon loss of *PRDM16* in the human organoids; 4. The gene is dysregulated upon loss of *Lhx2* in the mouse. This filtering resulted in the identification of 34 genes. These 34 genes are involved in Wnt signaling, cell adhesion, and ECM receptor interaction, offering a shared gene regulatory network involving the cooperation of PRDM16 and LHX2 **(Supplementary Table S6)**.

## DISCUSSION

PRDM16, a transcriptional regulator with reported histone methyl-transferase activity, has been identified as a critical factor regulating human adipose tissue development, hematopoietic development and is associated with cardiomyopathies [1,19,20]. Here we report a patient with haploinsufficiency in *PRDM16* showing severe neurological defects **(Fig. 1 and Supplementary** Fig. 1). Furthermore, *PRDM16* has been postulated to contribute to the 1p36 deletion syndrome [21,29]. Using a brain organoid-based approach, we incorporated the patient’s mutation to investigate and gain mechanistic insights into the functions of PRDM16 in human brain development **(Fig. 2-3, Supplementary** Fig. 1). In our organoid model, we observed diverse impacts of PRDM16 on human neurogenesis. Previous studies have identified 5 core genes *SKI, GABRD, RERE, PRDM16*, and *KCNAB2* responsible for most of the developmental defects observed in patients with 1p36 deletion syndrome. However, the contribution of PRDM16 to human neural development was mostly unknown. Our study indicates that the deletion of one copy of PRDM16 plays a primary role in the observed neurodevelopmental defects of patients with 1p36 syndrome. Additionally, specific findings with a transcriptomic and histological basis of organoids align with those from the mouse model.

## PRDM16’s role in embryonic progenitors

Our analysis of mutant organoids containing a mutation detected in a human patient uncovers aspects of PRDM16 function that are conserved across mice and humans. Our organoid data are consistent with previous findings that, in the embryonic mouse cortex, the absence of *Prdm16* decreases the number and cycling of Tbr2 progenitors and increases cell cycle exit. It is also consistent with a recent paper suggesting that the absence of *Prdm16* leads to the persistence of RG into late postnatal stages [16]. The increase was observed only in apical progenitors but not in the basal progenitors. In both the mouse-developing brain [41] and the human organoid systems, the loss of PRDM16 resulted in a decrease of basal or outer radial cells that are HOPX+ suggesting an evolutionary conserved role of PRDM16 in the expansion of apical radial glia. In addition, our results suggest that PRDM16 levels affect the cell cycle of the progenitors, where, in the case of the homozygous mutation, we observed increased proliferation, possible S-phase elongation, and an increase in the population of cells that exit the cell cycle. These results are consistent with those across mammals in which PRDM16 emerges as a factor that controls multiple aspects of the biology of progenitor cells [22,23,39,41]. A noteworthy unaddressed aspect in the current study pertains to an observation made by one of the researchers (J-M.B., unpublished data, supplementary video 1). This observation revealed that a subset of the adult *Prdm16* knockout mice exhibited mild or conspicuous epileptic seizures, mirroring a similar occurrence observed in the described human patient and in 1p36 deletion syndrome patients.

## PRDM16 affects cortical neurogenesis via WNT signaling and cell adhesion

By analyzing transcriptomic differences during cortical neurogenesis in the organoids, we found that the identity of the DEGs is consistent with previously reported roles for PRDM16 across species. Our results suggest that PRDM16 suppresses the proliferation of RG cells partly by inhibiting the canonical Wnt-signaling pathway. An increase in canonical WNT signaling in the cortex is associated with increased cell cycle exit of intermediate progenitors into neurons [56]. We postulate that decreased SVZ/O-SVZ progenitors observed following the loss of PRDM16 may result from this increase in WNT signaling [56]. In our organoids, we discovered that the loss of PRDM16 disrupts the delicate balance of WNT signaling, potentially affecting cell proliferation and adhesion processes.

## PRDM16 controls multiple aspects of corticogenesis across mice and humans

Loss of PRDM16 in mice is associated with defects in cell migration that result in the formation of neuronal heterotopias that persist into adulthood [16,41]. Interestingly, neuronal heterotopias is a common cause of seizures in humans [57,58]. Therefore, the presence of severe seizures in the patient with PRDM16 haploinsufficiency described in this study opens the possibility that PRDM16 also regulates neuronal migration in the human brain. Histological and transcriptomic analysis of human organoids suggests a PRDM16 control of various aspects of neural stem cell biology including progenitor proliferation, transition of apical PAX6+ RG progression to basal HOPX+ RG, aspects of cell adhesion, and migration. This is consistent with previous findings in mice showing that PRDM16 promotes cortical RG lineage progression into ependymal cells or adult neural stem cells [15,16]. Future experiments with more sophisticated human organoids systems that allow the study of PRDM16’s role in the generation of basal RG, neuronal migration that ultimately contribute to the expansion of the human brain will be necessary for addressing this question.

## An evolutionarily conserved PRDM16-related gene regulatory network

We found that thousands of PRDM16 binding sites across the genome of cortical RG have been conserved since the divergence of humans and mice around 90 million years ago. This result highlights the importance of PRDM16-mediated transcriptional regulation in the development and evolution of the human brain. Furthermore, whereas mice haploinsufficient for *Prdm16* are grossly normal, the loss of one copy of *PRDM16* in humans had strong deleterious effects in cortical RG in vitro and in a human patient, suggesting a more critical function of PRDM16 in human brain development in comparison to rodents. Our analysis of conserved PRDM16 binding sites between mouse and humans indicates that regulation of developmental enhancers is likely the primary mechanism by which PRDM16 controls gene transcription in the developing human brain, similarly to what was found in the mouse cortex (40). Future studies may be directed at identifying human-specific PRDM16-bound enhancers and their potential role in the evolution of human-specific brain features, such as an increase in the number of basal RG cells.

An outstanding open question is how PRDM16, a regulator that is expressed in multiple somatic tissues, controls neural-specific transcriptional programs. Our motif analyses within conserved PRDM16 binding sites in the human brain revealed an enrichment of neurogenesis-related factors. Although we did not detect a PRDM16 motif enrichment, a PRDM motif enrichment in the data was found (PRDM9 and PRDM14). The fact that a PRDM16 motif was not found may stem from the fact that the data for motif discovery came from PRDM14 and PRDM9 ChIP-seq datasets. Of note, neither PRDM9 nor PRDM14 are expressed in the mouse cortex and the developing human cortex (www.brainspan.org). Therefore, the PRDM9 and PRDM14 motif discovery suggest that PRDM16 could be binding directly to some sequences, albeit relatively few, in the human cortex, which is in contrast to what we found in the mouse cortex.

Importantly, the LHX2 DNA binding motif was the most enriched within conserved PRDM16-bound regions. LHX2 is a pleiotropic transcription factor whose loss of function phenotypes in the mouse are well characterized [51]. We found that loss of PRDM16 in human organoids resulted in a dysregulated GRN similar to the one observed in *Prdm16* and *Lhx2* cKO mice. Moreover, we show that PRDM16 and LHX2 bind highly overlapping genomic regions near neurogenic genes in the mouse cortex (Figure 6F). Loss of either transcription factor upregulated neurogenesis, but its effect on progenitor identity was different in nature; while loss of PRDM16 leads to increased PAX6+ cells and prolonged radial glial lifetime in the mouse brain [16], loss of LHX2 in the mouse leads to reduced *Pax6* expression and to a reduced number of PAX6-positive cells [54]. This discrepancy might result from the expression of genes differentially regulated by PRDM16 and LHX2. However, we found that both, PRDM16 and LHX2, may be involved in governing partially overlapping progenitor cell cycle exit [53,54]. Our findings in the organoid system also suggest an increased canonical WNT signaling in mutant organoids (Fig. 4), which suggests a repressive role for PRDM16. LHX2 and CTNNB1 act cooperatively to balance cortical progenitor proliferation and differentiation [53]. We postulate that PRDM16 cooperates with LHX2 to inhibit the canonical Wnt signaling pathway in the cortex. More experiments are needed to determine whether PRDM16 and LHX2 form a transcriptional complex in RG and whether LHX2 is necessary for the recruitment of PRDM16 to a subset of enhancer regions in the cortex. Our results uncovered a conserved functional interaction between PRDM16 and LHX2 in neural stem cells that governs brain development across species.

## Author Contributions

### Methods

#### Ethics statement

Work with hESC (WIBR3, NIHhESC-10-0079) and genome editing was carried out with approval from the Weizmann Institute of Science IRB (Institutional Review Board). The information from the patient and the use of blood samples was consented to by the parents and approved by the IRB in Sheba Med Ctr.

#### hESCs and brain organoid cultures

WIBR3 (NIHhESC-10-0079) hESCs and isogenic PRDM16 mutant lines were grown and maintained on irradiated MEFs in optimal naive NHSM conditions (RSET-Stem Cell Technologies INC, [59]).

#### CRISPR/Cas9 genome editing

The patient’s mutation was introduced in hESC (WIBR3, NIHhESC-10-0079) using hSpCas9n nickase and chimeric guide RNA (gRNA), and single strand oligonucleotide (ssOD) repair.

The sequences of the gRNA pair are: 5’-GGCCGGGCGGCACTGTGCCC; 5’- GTAAGACCCCTCCCCCAAAC. The ssOD sequence is 5’- TTAAGGACATTGAGCCAGGTGAGGAGCTGCTGGTGCACGTGAAGGAAGGCGTCTACCCCCGGGCACAGTGCCGCCCGGCCTGGACGGTAAGACCCCTCCCCCAAACCGGGCCACGGCCCC.

Three days after transfection, the cells were subjected to FACS and plated at a density of 2,000 cells per 10 cm plate on irradiated MEFs, allowing for the growth of single-cell-derived colonies. The mutation was confirmed by PCR, restriction enzyme digestion, and Sanger DNA sequencing. In this study we used clones #62 and #121, which are heterozygous, and clone #95 which is homozygous.

#### Visualization and quantification of gene expression in single cells

The data for UMAP, expression levels, and phase annotation of the pallial excitatory neuron lineage were obtained from [45]. These cells were computationally isolated by selecting *EMX1*-expressing clusters from samples that were dissected from the human telencephalon 5-14 PCW.

To test for differential gene expression of *AURKA* and *PRDM16* in radial glia between progenitor states, we focused on *EMX1*-expressing clusters from 11-14 PCW. All of these samples were prepared using the 10X V3 chromium chemistry, which enabled us to compare their gene expression patterns. To determine statistical significance, we employed a two-sided Mann-Whitney test with Bonferroni correction.

#### Cortical organoids

Cortical organoids were grown as previously described [46]. Briefly, about 9000 ES cells per well were dispensed into low adhesive v-shaped 96-well plates, and aggregates were formed. Forebrain organoids media 1 **(Table 2)**, containing WNT inhibitor and TGF inhibitor, was exchanged every other day. On day 18, organoids were transferred to a 10-cm non-cell adhesive plate and cultured in a second forebrain media **(Table 3)** in suspension. Organoids were collected after two weeks, on day 32.

**Table 1:**
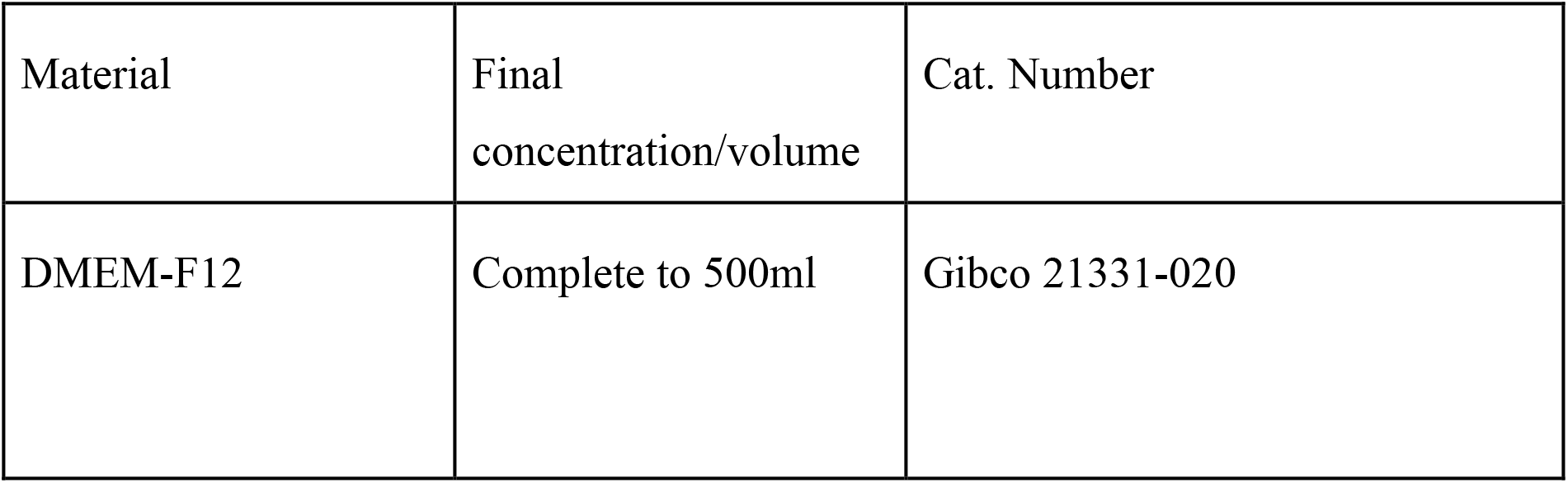
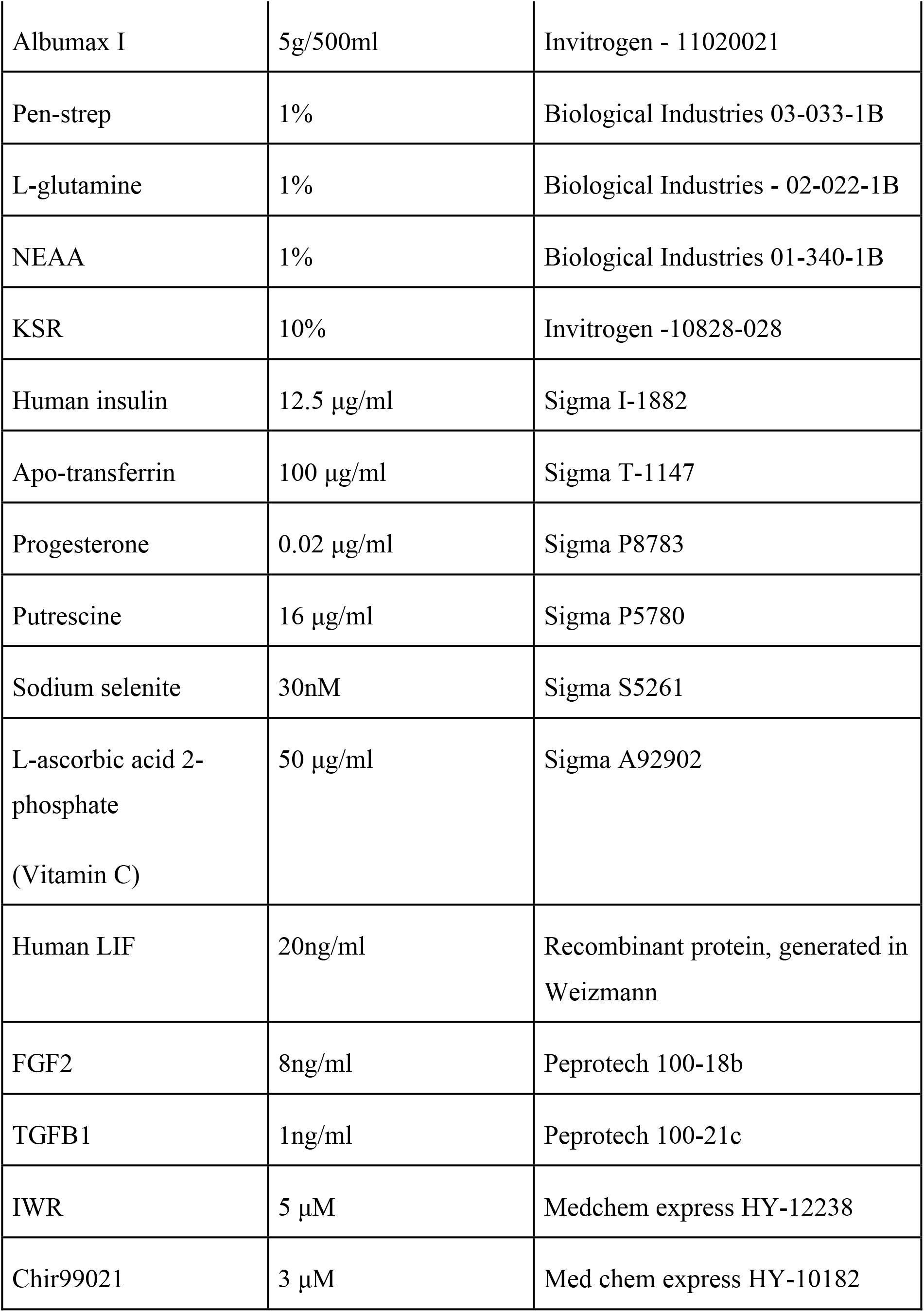
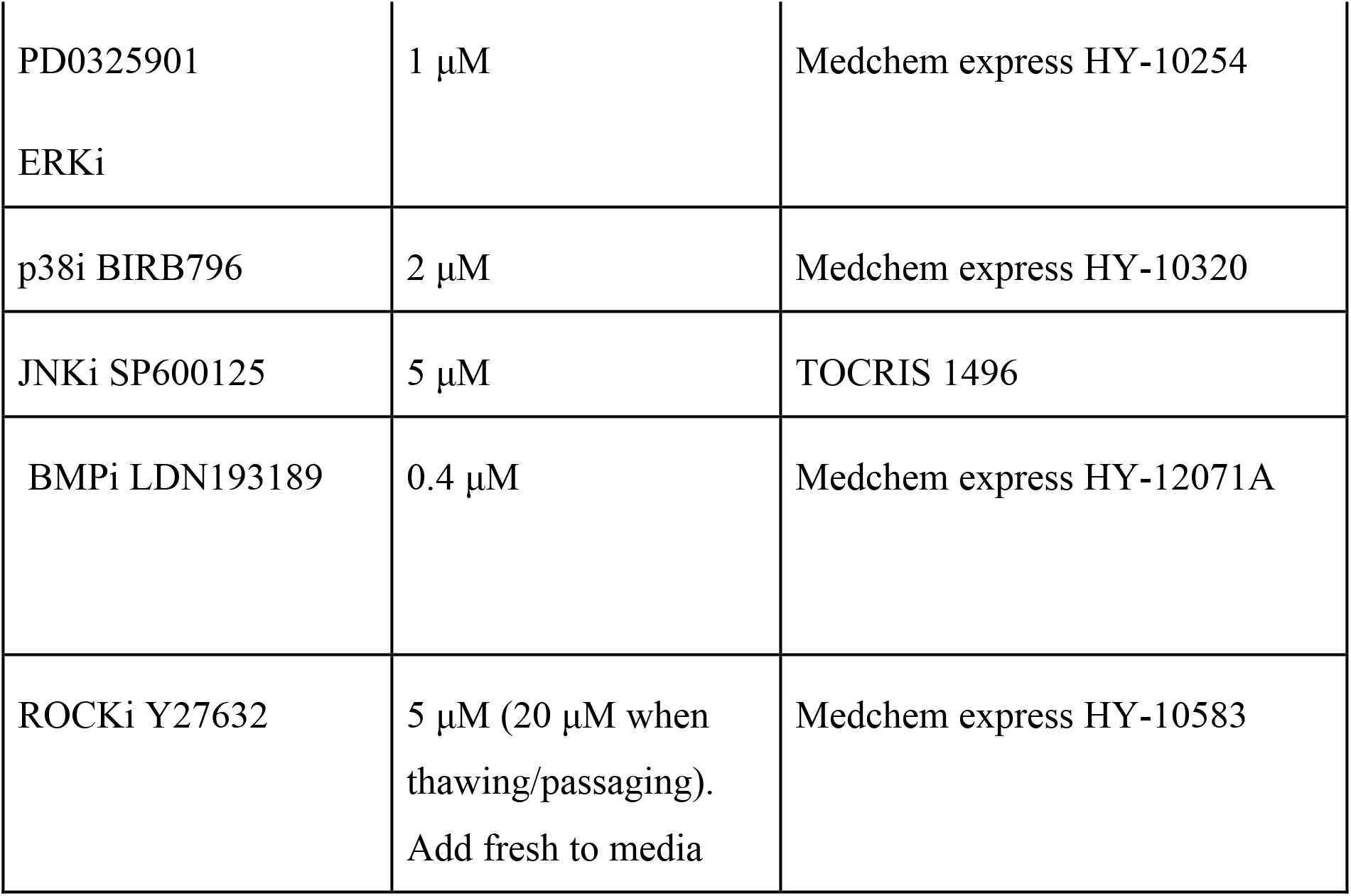
ESCs maintained in Naïve media (NHSM media)

**Table 2:** Forebrain organoids media 1 (days 0-17)

|  | Material | Final Concentration | Catalog Number |
| --- | --- | --- | --- |
| 1 | DMEM | Complete to 100% | Gibco 41965-039 |
| 2 | KSR | 20% | Invitrogen 10828-028 |
| 3 | NEAA | 1% | Biological Industries 01-340-1B |
| 4 | Pen-strep | 1% | Biological Industries 03-033-1B |
| 5 | Sodium Pyruvate | 1% | Biological Industries 03-042-1B |
| 6 | 2-beta<br>mercaptoethanol | 0.1 mM | Gibco 3150-10 |
| 6 | TGFBi (SB431542) | 5mM | Medchem express HY-10431 |
| 7 | IWR | 3mM | Medchem express HY-12238 |
| 8 | ROCKi Y27632<br>(until day 6) | 20mM | Medchem express HY-10583 |

**Table 3:** Forebrain organoids media 2 (days 18-34)

|  | Material | Final Concentration | Catalog Number |
| --- | --- | --- | --- |
| 1 | DMEM/F12 | Complete to 100% | Gibco 21331-020 |
| 2 | Glutamax | 1% | Gibco 35050-038 |
| 3 | N2 Supplement | 1% | Gibco 17502048 |
| 4 | Chemically Defined<br>Lipid Concentrate | 1% | Gibco 11905-031 |
| 5 | Pen-strep | 1% | Biological Industries 03-033-1B |

#### RNA-seq of organoids

Total RNA was extracted using the miRNeasy Mini kit (Qiagen, Hilden, Germany). RNA concentration and integrity were measured using Nanodrop (Thermo Scientific) and an Agilent Tapestation 4200. Total RNAseq libraries from 1 µg of ribosomal depleted RNA using TruSeq Stranded Total RNA Library Prep (Illumina). These libraries were sequenced on the Illumina NOVA-seq platform to obtain 150 bp paired-end libraries. Sequencing libraries were prepared from 1000 ng of purified total RNA pooled for biological replicates of 5-8 organoids.

#### RNA-seq analysis of organoids

RNA-seq reads were processed using the UTAP pipeline [60] and were previously described in [61]. Briefly, fastq files were aligned to the GRCm38 reference genome using STAR (version v2.4.2a). Gene read count was performed using the qCount function from the QuasR Bioconductor package (v.1.34) with default parameters. Differentially expressed genes were identified with the DESeq2 Bioconductor package (v.1.34). Genes with log2foldchange ≥ 1 and ≤ −1 with padj ≤ 0.05 and baseMean ≥ 10 were considered differentially expressed. Clustering was performed with the kmeans function in R. Differentially expressed genes were analyzed using ShinyGO [50].

#### qRT-PCR

Total RNA was extracted from a pool of 5-8 organoids using the RNeasy Mini kit (Qiagen) under the manufacturer’s protocols. RNA was converted to cDNA by qScript cDNA Synthesis Kit (quantabio). Real-time PCR with SYBR FAST ABI qPCR kit (Kapa Biosystems) was performed using QuantStudio 5 Real-Time PCR System (Applied Biosystems). Each group contained three biological repeats, each with three technical repeats. If one of the technical repeats differs from the other two in more than one cycle, it is removed from the calculation. Thus, in each statistical group, n<=9. The delta CT values were used for statistical analysis, with RPS29 as an internal control.

The fold change was calculated using the delta-delta CT (ddCT) method, in which each delta CT value was normalized to the average CT value of the control line at the specific time point. Fold change=2^-ddCT.

Primers were taken from the primerBank database and are detailed in **Table 4**

**Table 4:**
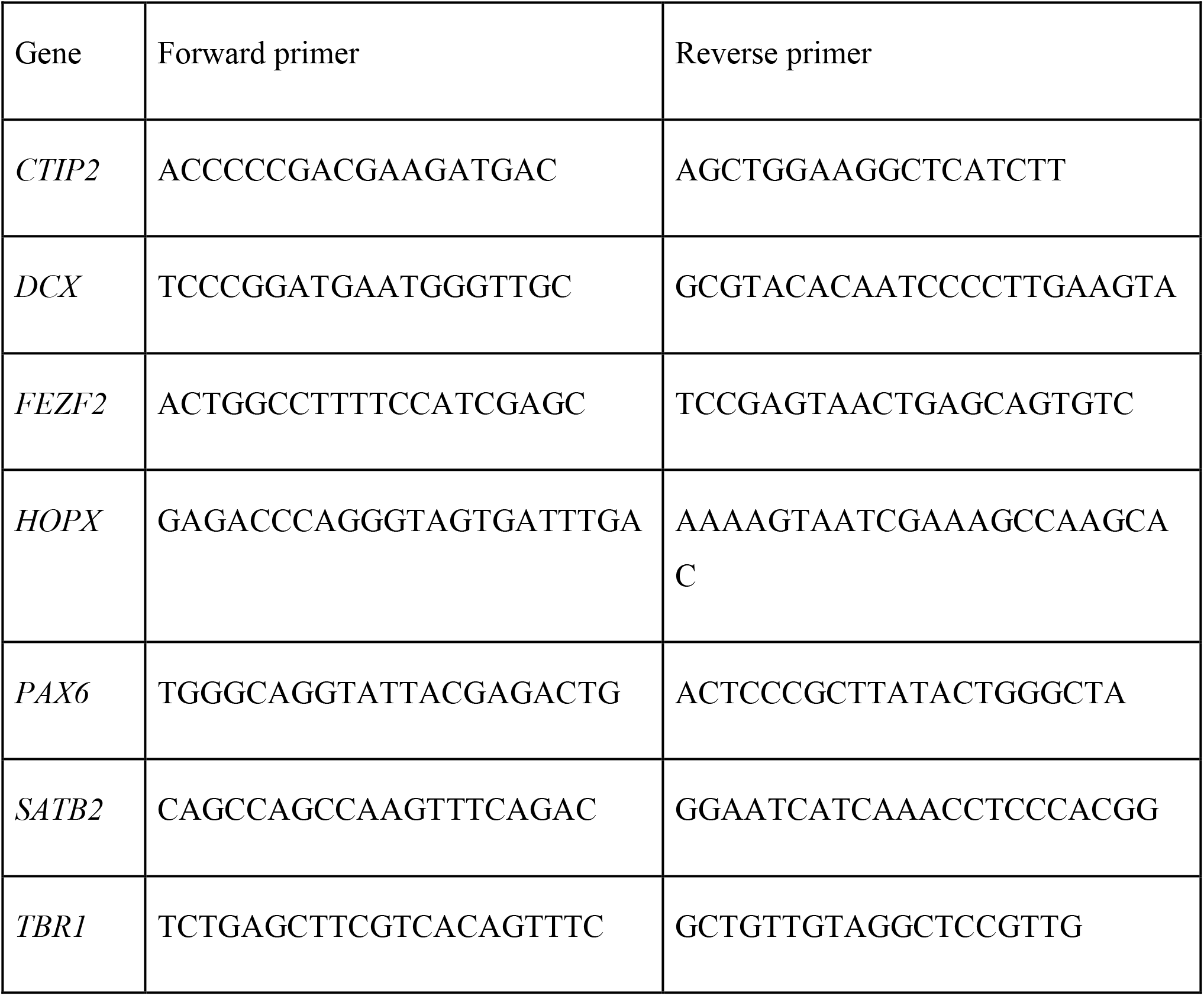
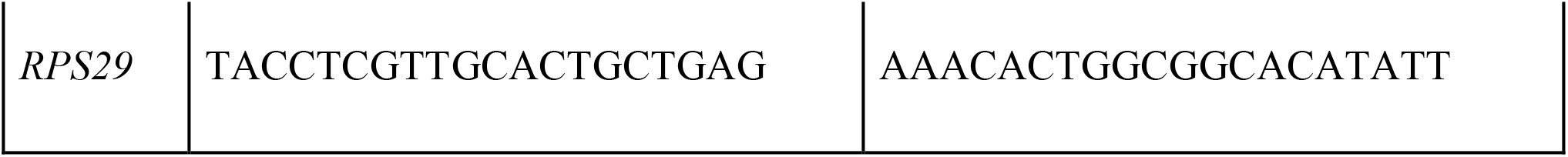
qPCR primers.

**Table 5:** List of antibodies used for this experiment.

| **Antibody** | **Host** | **Clonality** | **Dilution** | **Company** | **Catalogue Number** |
| --- | --- | --- | --- | --- | --- |
| KI67 | Mouse | Monoclonal | 1:400 | BD Pharmingen | 550609 |
| pHH3 | Rabbit | Polyclonal | 1:200 | MilliporeSigma | 06-570 |
| cleaved Caspase3 | Rabbit | Polyclonal | 1:200 | Cell signaling | 9661 |
| SOX2 | Mouse | Monoclonal | 1:100 | SCBT | sc-365823 |
| PAX6 | Rabbit | Polyclonal | 1:100 | BioLegend | 901301 |
| HOPX | Rabbit | Polyclonal | 1:200 | MilliporeSigma | HPA030180 |
| MAP2 | Mouse | Monoclonal | 1:200 | Sigma-Aldrich | M9942 |
| CTIP2 | Rabbit | Polyclonal | 1:200 | Abcam | ab28448 |
| Trb1 | Rabbit | Polyclonal | 1:100 | Abcam | ab31940 |
| β-catenin | Mouse | Monoclonal | 1:200 | BD Biosciences | 610153 |
| LEF1 | Rabbit | Monoclonal | 1:200 | Cell signaling | 2230 |
| N-cadherin | Mouse | Monoclonal | 1:50 | DSHB | 6B3 |
| Collagen Type I | Goat | Polyclonal | 1:200 | SouthernBiotech | 1310-01 |

#### Organoid Histology

Immunohistochemistry was performed on day 32 cortical organoids from both control and *PRDM16* mutant cell lines. EdU in DMSO was incorporated into the organoids at a working concentration of 10 µM in culture media for 30 minutes at 37⁰C. The organoids were then fixed in 4% paraformaldehyde for 3-4 hrs at 4⁰C on a shaker and later stored overnight in 20% sucrose at 4⁰C on a shaker. For the cryosection, OCT blocks were prepared, and 12 µm slices were collected on the positively charged microscope slides such that each sectioned organoid was represented three times on the same slide at three different heights of the organoid in succession. Sections from both control and mutant organoids were collected on the same slide. After air drying, the microscope slides for overnight; for IHC, antigen retrieval was performed by boiling the sections at 90-95⁰C in 10 mM sodium citrate buffer (pH 6) for 5-6 min. Blocking was performed for 1 hr using 10 % horse serum + 10 % FBS + 0.1 % Triton-X 100 in 1X PBS. For EdU detection, Click-iT^®^ EdU Flow Cytometry Assay kit was used. The slides were incubated with the reaction mixture comprising of 100 mM Tris (pH 8.5) + 1 mM CuSO4 + 100 mM ascorbic acid + 2.5 µM Cy5 azide in 1X PBS for 30 minutes at room temperature. Primary antibody incubation was performed overnight at 4⁰C, followed by incubation with secondary antibodies conjugated with Alexa Fluor^®^ 488 and Alexa Fluor^®^ 555 at room temperature for 2 hours. Counter-nuclear staining was performed with Hoechst 33352 (1:2000 dilution in 1X PBS). The primary antibodies were used in the following dilutions: KI67 (mouse, 1:400, BD Pharmingen), pHH3 (rabbit, 1:200, MilliporeSigma), cleaved Caspase3 (rabbit, 1:200, Cell Signaling), SOX2 (mouse, 1:100, SCBT), PAX6 (rabbit, 1:100, BioLegend), HOPX (rabbit, 1:100, MilliporeSigma), MAP2 (mouse, 1:200, Sigma-Aldrich), CTIP2 (rabbit, 1:200, Abcam), TBR1 (rabbit, 1:100, Abcam), β-catenin (mouse, 1:200, BD Biosciences), LEF1 (rabbit, 1:200, Cell Signaling), N-cadherin (mouse, 1:50, DSHB) and Collagen Type I (goat, 1:200, SouthernBiotech). After the slides with mounting media (Prolong^TM^ Gold Antifade reagent – Invitrogen by Thermo Fischer Scientific, REF – P36930) had dried completely, images were captured with Oxford Dragonfly confocal microscope at 25X and stitched to get complete images of the whole organoid sections using the Fusion Shell software. Higher-resolution images were captured with an oil-immersion lens at 63X.

The images were then analyzed using the Spots Analysis of the Oxford Imaris software to determine the percentage of the total number of cells that expressed a particular cell-type specific marker and compare this parameter between control and mutant organoids. The total number of cells is considered the same as the total number of Hoechst^+^ nuclei. Furthermore, the number of cells expressing markers like KI67, pHH3, EdU, SOX2, PAX6, HOPX, CTIP2, TBR1, and LEF1 that have nuclear staining, were determined by setting up a colocalization filter under the Spots analysis with a suitable threshold for the distance between Hoechst and other fluorescent marker spots in order to avoid any nonspecific signal. The germinal area was determined by the average area of rosette-like arrangement of cells with the organoid sections. These rosettes have a typical neuroepithelium-like arrangement of progenitor cells. For extra-nuclear markers like cleaved Caspase3, MAP2 and Collagen Type I, the percentage of total area of the organoid section that has signal from these markers were determined by setting up a suitable intensity threshold in Fiji ImageJ software. For markers like β-catenin and N-cadherin, mean intensity and intensity density of the signal in the total area of the organoid section was calculated in Fiji ImageJ software to compare the levels of expression of the respective markers between control and mutant organoids. These values were then compared using statistical tests in GraphPad Prism to determine statistically significant differences, if any.

#### Human samples

Research performed on samples of human origin was conducted according to protocols approved by the institutional review boards (IRB) of Beth Israel Deaconess Medical Center. Fetal brain tissue was received after release from clinical pathology, with a maximum post-mortem interval of 4 h. Cases with known anomalies were excluded. Brain tissue was collected from 15 and 22 gestation weeks (GW). Tissue was transported in HBSS medium on ice to the laboratory for research processing.

#### Human Sample Collection and Crosslinking

Postmortem human fetal samples of the cerebral cortex were obtained at GW15 and GW22. The dissected cortical regions included the VZ and outer SVZ, which is the tissue that contains most of the PRDM16^+^ cells. Tissue samples were either partially fragmented with papain and frozen in cryo-preservation media (GW15) or immediately frozen in liquid nitrogen (GW22) and kept at −80° C. To collect the frozen samples, the fragmented tissue was resuspended in Hibernate E medium (Thermo Fisher Scientific) and allowed to sediment for 5 min. Hibernate E was pipetted out, and the tissue was crosslinked as indicated below. For processing freshly frozen samples of the unfragmented human cortex, the tissue was ground inside a Covaris tissue TUBE (Covaris 520001) on top of dry ice with alternating rounds of submerging the tube in liquid nitrogen to avoid thawing of the sample until a fine tissue powder was generated, as described previously (Savic D. et al. 2013. Epigenetics and Chromatin). All human samples were dual-crosslinked by a serial treatment with 1.5 mM glycol bis[succinimidyl succinate] (EGS, Thermo Fisher Scientific) in PBS1X for 20 min at room temperature (RT) with rotation, followed by treatment with 1% ethanol-free Paraformaldehyde (PFA, Electron Microscopy Sciences) for an additional 10 min at RT. Crosslinking was quenched by adding glycine to a final concentration of 125 mM and rotating for 5 min at RT. Fixed samples were washed with 10 ml of cold PBS 1X plus protease inhibitor. The tissue was centrifuged at 1500 g for 6-8 min, and the PBS wash was repeated. The PBS was removed as much as possible after the second wash, and the pellets were either flash-frozen in liquid nitrogen and stored at −80° C or immediately used for chromatin immunoprecipitation.

#### Human Chromatin Immunoprecipitation and Sequencing (ChIP-seq)

The procedure for ChIP-seq was done as previously described [41], with some minor modifications. Briefly, crosslinked tissue was thawed, resuspended in cell lysis buffer (20 mM Tris-HCl pH 8.0, 85 mM KCl, 0.5% NP40), and incubated for 30 min to 1 hr on ice with vortexing every 5-10 min. Nuclei were pelleted at 1500 g for 5 min at 4° C, resuspended in SDS buffer (0.2% SDS, 20 mM Tris-HCl pH 8.0, 1 mM EDTA), and sonicated in a Covaris S2 ultrasonicator using the following settings: Intensity = 5, duty cycle = 10%, cycles per burst = 200, treatment time = 13 min. The sonication step eliminated the residual nuclei clumps. Nuclei were pelleted at 18,000 g for 10 min and resuspended in 2X ChIP dilution buffer (0.1% sodium deoxycholate, 2% Triton X-100, 2 mM EDTA, 30 mM Tris-HCl pH 8.0, 300 mM NaCl). At this point, around 2.5% of the sample volume was set aside for later use as input control. The remaining sample was incubated overnight at 4° C with rotation with 5 mg of sheep anti-PRDM16 antibody (R&D Systems, AF6295) or 5 mg of sheep IgG control antibody. The next day, 40 ml of protein G magnetic beads (Novex) were added to the chromatin and incubated at 4° C with rotation for 2 hrs. The beads were pooled with a magnet, the supernatant was completely removed, and the beads were washed twice for 10 min with low salt wash buffer (0.1% SDS, 1%Triton X-100, 2 mM EDTA, 20 mM Tris-HCl pH 8.0, 150 mM NaCl), high salt wash buffer (0.1% SDS, 1% Triton X-100, 2 mM EDTA, 20 mM Tris-HCl pH 8.0, 500 mM NaCl), LiCl wash buffer (0.25 M LiCl, 0.5% NP40, 0.5% sodium deoxycholate, 1 mM EDTA, 10 mM Tris-HCl pH 8.0), and TE buffer (10 mM Tris-HCl pH 8.0, 1 mM EDTA). The chromatin was eluted from the beads in 90 ml of freshly prepared ChIP elution buffer (1% SDS, 0.1 M NaHCO3) at 65° C for 30 min with rotation. The samples were incubated at 65° C overnight in reverse crosslinking solution (250 mM Tris-HCl pH6.5, 62.5 mM EDTA pH 8.0, 1.25 M NaCl, 5 mg/ml of Proteinase K) followed by extraction of the genomic DNA with phenol/chloroform/isoamyl alcohol. The DNA was precipitated with 1/10 volume of 3M sodium acetate pH 5.0, glycogen, and two volumes of ethanol and resuspended in TE low EDTA (10 mM Tris-HCl pH 8.0, 0.1 mM EDTA). All samples were treated with RNase A (100 mg/ml) for 30 min to 1 hr at 37° C. We used the Ovation Ultralow System V2 kit (NuGEN) for library preparation according to the manufacturer’s instructions. The libraries were size-selected in the 100-800 bp range using agarose gel electrophoresis and DNA gel extraction (QIAGEN). Libraries were sequenced in an Illumina HiSeq 2500 sequencer to a sequencing depth of 40-50 million reads per sample. Human Histology and immunofluorescence were performed as previously described in [41].

#### RNA-seq & ChIP-seq overlap

To identify direct Targets of PRDM16 and LHX2 we associated the peaks with the nearest gene using HOMER and compared it with the RNA-seq. Overlap of RNA-seq and ChIP-seq genes were subjected to hypergeometric statistical analysis using http://nemates.org/MA/progs/overlap_stats_prog.html.

#### Mice

All animal protocols were approved by the Institutional Animal Ethics Committee of the Tata Institute of Fundamental Research, Mumbai, India. The floxed LIM homeobox2 (*Lhx2*) line (*Lhx2lox/lox*) and *Emx1CreYL* lines used in this study have been described previously by [62,63]. The *Emx1CreYL* [63] was obtained as a gift from Prof. Yuqing Li at the University of Florida College of Medicine. The floxed *Lhx2* line was a gift from Prof. Edwin Monuki at the University of California, Irvine. Timed pregnant female mice were obtained from the Tata Institute animal breeding facility, and embryos of both sexes were used for the experiments, with the *Emx1CreYL* contributed from the male parent. Noon of the day the vaginal plug was observed was considered E0.5. Early-age embryos were staged by somite number, genotyped using PCR and assigned to groups accordingly. Controls used for each experiment were age-matched littermates. The mT/mG reporter mouse line was obtained from JAX labs Stock No. 007576; this reporter was used to check for cre activity in the brain. All animals were kept at an ambient temperature and humidity, with a 12 hr. light-dark cycle and food available ad libitum. Primers used for genotyping were: Cre F: 5′ATTTGCCTGCATTACCGGTC3′, Cre R: 5′ATCAACGTTTTCTTTTCGG3′, Cre-positive DNA shows a band at 350 bp. *Lhx2* cKO forward: 5’ACCGGTGGAGGAAGACTTTT3’, *Lhx2* cKO reverse: 5’CAGCGGTTAAGTATTGGGACA3’. The band sizes for this PCR are as follows: Wild-type: 144 bp, *Lhx2Cko*: 188 bp.

#### Mouse ChIP-seq and data analysis

LHX2 ChIP-seq was performed in 4 biological replicates of each sample E12.5 Neocortex. Input DNA was used as a control and sequenced with the respective samples for all ChIP-seq experiments. The mouse PRDM16 ChIP-seq PEAKS was obtained from [41]

##### Tissue processing

Neocortical and hippocampal tissue were dissected from E12.5 Swiss mice and collected in ice-cold PBS containing 0.5% glucose and a protease inhibitor cocktail (P8340). Tissue was crosslinked using 1% formaldehyde (#47608) for 8 minutes, followed by quenching with 125 mM glycine for 5 minutes at RT. The chromatin was sheared using a focused sonicator (Covaris) to obtain fragments of 100-300 bp. 100 µg (for LHX2) or of sheared chromatin was used to set up an IP with the LHX2 antibody from Santa Cruz Biotechnology (Sc-19344) and 10% of the chromatin volume was stored as input. Dynabeads A and G were mixed in a 1:1 ratio and used to pull down the antibody-protein complex. Beads were washed 3 times with low salt buffer (20 mM Tris HCl pH 8.0, 150 mM NaCl, 2 mM EDTA, 0.1% SDS, 1% Triton X-100), followed by 2 washes with high salt buffer (20 mM Tris HCl pH 8.0, 200 mM NaCl, 2 mM EDTA, 0.1% SDS, 1% Triton X-100), 1 wash with LiCl buffer (0.25 M LiCl, 1 mM EDTA, 10 mM Tris HCl pH 8.0, 1% NP-40, 1% sodium deoxycholate) and 2 washes with TE buffer (10 mM Tris HCl pH 8.0, 1 mM EDTA). The beads were resuspended in 150 µl of elution buffer (0.1M NaHCO3, 1% SDS) and at 65°C for 30 minutes at 1000 rpm. The eluate was collected in fresh tubes and the elution was repeated to obtain a total eluate of 300 µl. The IP and input samples were reverse cross-linked using 20 µL of 5 M NaCl and 2 µL of RNAseA (10 mg/ml), and incubated overnight at 65°C at 800 rpm. The samples were then treated with 20 µL of 1 M Tris pH 8.0, 10 µL of 0.5 M EDTA and 2 µL of Proteinase K (20 mg/ml) and incubated at 42°C for 1 hr at 800 rpm. Samples were purified using phenol: chloroform: isoamyl alcohol and DNA was precipitated at −20°C using 2X volume of 100% ethanol, 100 mM sodium acetate and Glycoblue (#AM9515). DNA pellets were resuspended in nuclease-free water and quantified using a Qubit fluorometer (Thermo Fisher Scientific, USA) for downstream processing.

##### Library preparation, sequencing and data analysis

An equal amount of DNA (∼5-8 ng) was used as an input for library preparation and libraries were prepared using an NEB Ultra II DNA library prep kit (NEB, USA). Sequencing reads (100 bp PE) were obtained on the HiseqX platform at Macrogen, Korea.

Sequencing reads were trimmed using TrimmomaticPE for Truseq2:PE adapters and were aligned to the mouse mm10 genome using the default parameters of BWA. Aligned reads were subsampled to 25 million reads for each sample using BBMap. For the LHX2 ChIP-seq the QC, peak calling was performed using default parameters in PePr [64]. Only statistically significant peaks were used for further analysis (p-value 0.0001 and fold change over input: cut off >10 fold).

#### Mouse RNA sequencing, library preparation, and analysis

The mouse Neo-cortices were dissected from E12.5 control and *Lhx2* mutant brains and stored in TrizolⓇ reagent. Tissue dissected from 4 embryos was pooled to obtain 5 µg RNA for each of the 3 biological replicates. After library preparation, sequencing was performed on the Illumina platform to achieve 150 bp reads to generate 30 Million paired-end reads per sample. FastQC was performed as described in (https://www.bioinformatics.babraham.ac.uk/projects/fastqc/), and reads > 30 Phred scores were aligned using HISAT2 [65]. Feature counts were used to quantify the number of reads per transcript. Differential expression analysis was performed using EdgeR [66] on the R platform (v3.4.0). Genes showing |log2 Fold change | ≥0.58 and FDR < 0.05 were used for further analysis. Gene ontology analysis was performed using WebGestalt or gShinyGO 0.76 [50,67]. Bar plots were created with GraphPad Prism V9.1.0. Gene-based heatmaps were plotted using normalized reads on Morpheus (Morpheus, https://software.broadinstitute.org/morpheus). Venn diagrams were initially generated using Venny 2.1.0 (https://bioinfogp.cnb.csic.es/tools/venny/) and edited using Adobe Photoshop (CC 2017).

## Acknowledgments

We thank Prof. Sten Linnarsson for generously sharing his data with us.

## Data Availability

The Organoid RNA-seq data related to this manuscript has been deposited to GEO under the submission ID : GSE239855

The mouse LHX2 ChIP-seq and related RNA-seq data are part of publicly available BioProjects: PRJNA882662, PRJNA883170

