## Supplementary Fig. for "PRDM16 co-operates with LHX2 to shape the human brain"

### SUPPLEMENTARY FIGURES

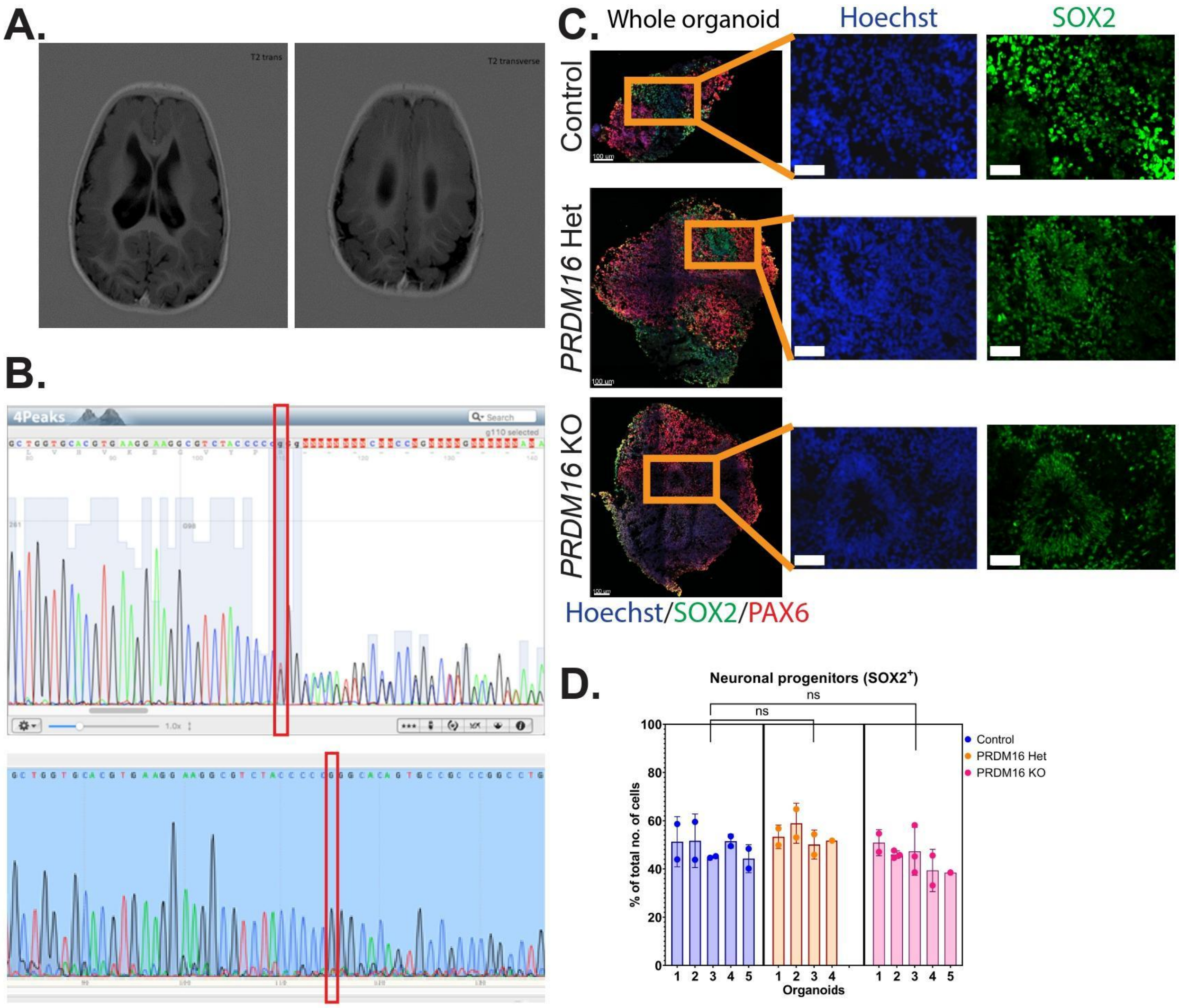


**Supplementary Fig. 1**

**A.** Additional MRI scans from the patient. **B.** Chromatograms of Sanger sequencing showing on the top the mutation in the patient's iPSCs (heterozygous), and on the bottom one of the homozygous hESCs PRDM16 mutated lines. **C.** Immunohistochemistry on 12 µm cryosections of day 32 control and *PRDM16* mutant cortical organoids showing the cortical progenitor markers – SOX2 and PAX6 (imaged using 25X objective lens, scale bar represents 50 µm in insets) **D.** Percentage of total number of cells that are $SOX2^+$ neuronal progenitors, in day 32 control and PRDM16 mutant cortical organoids (Nested One-Way ANOVA, n = 5, α = 0.05, ns: not significant; p-values: * <= 0.1; ** <= 0.01;

*** <=0.001; **** <=0.0001, each data point on the graph represents an image of the organoid along its z-axis)

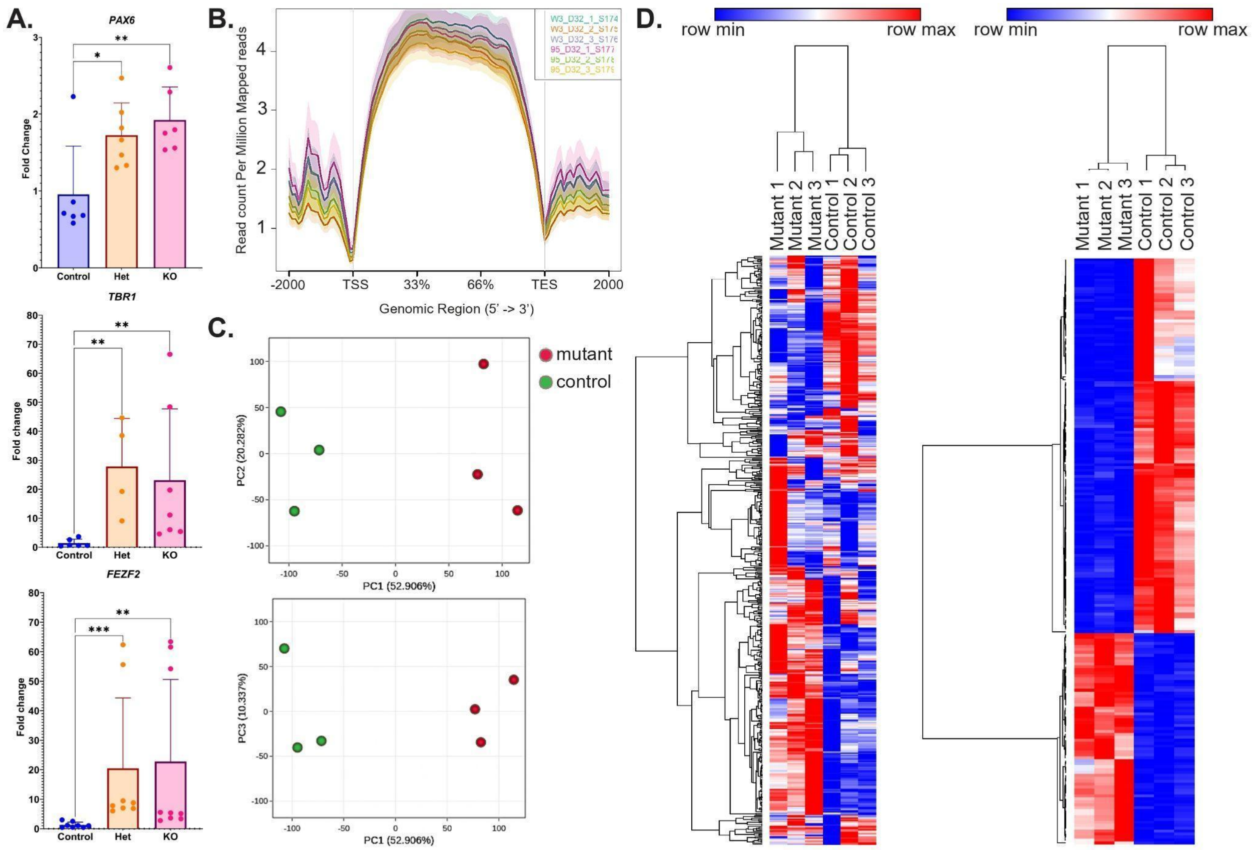


**Supplementary Fig. 2**

**A.** qPCR analysis derived from control, heterozygous, and mutant PRDM16 cortical organoids on day 32 revealed increased levels of *PAX6, TBR1* and *FEZF2* in heterozygous and homozygous mutants. **B.** Coverage plot of all RNA-seq samples. **C.** PCA plots of all RNA-seq samples. **D.** Heatmaps of all RNA-seq samples depict sample wise relationships of the Top 2000 expressed genes and top 200 DEGs identified by DESeq2. Each line represents a gene with normalized reads used as the input to generate this plot using Morpheus, https://software.broadinstitute.org/morpheus.

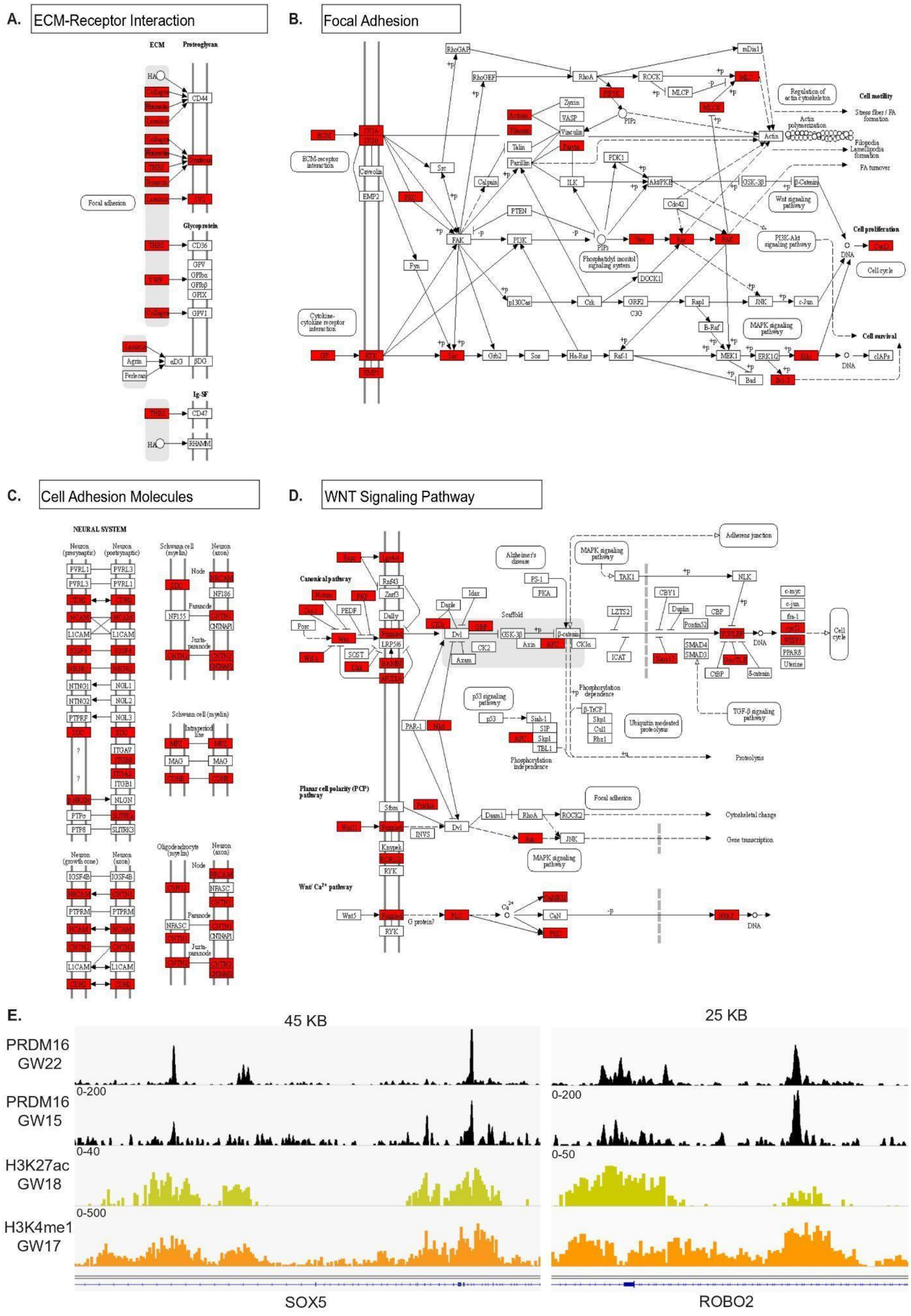

A.
ECM-Receptor Interaction
B.
Focal Adhesion
C.
Cell Adhesion Molecules
D.
WNT Signaling Pathway
E.
45 KB
25 KB
PRDM16 GW22
PRDM16 GW15
H3K27ac GW18
H3K4me1 GW17
0-200
0-40
0-500
0-50
SOX5
ROBO2

**Supplementary Fig. 3**

**A-D.** KEGG plot view of all DEGs (red) identified in the mutant versus control organoid RNA-seq data shows multiple genes affected in all 4 exemplified pathways. **E.** Genomic tracks displaying the overlap of PRDM16 binding sites with enhancer marks H3K27ac and H3K4me1 within the *SOX5* and *ROBO2* loci in the human cortex

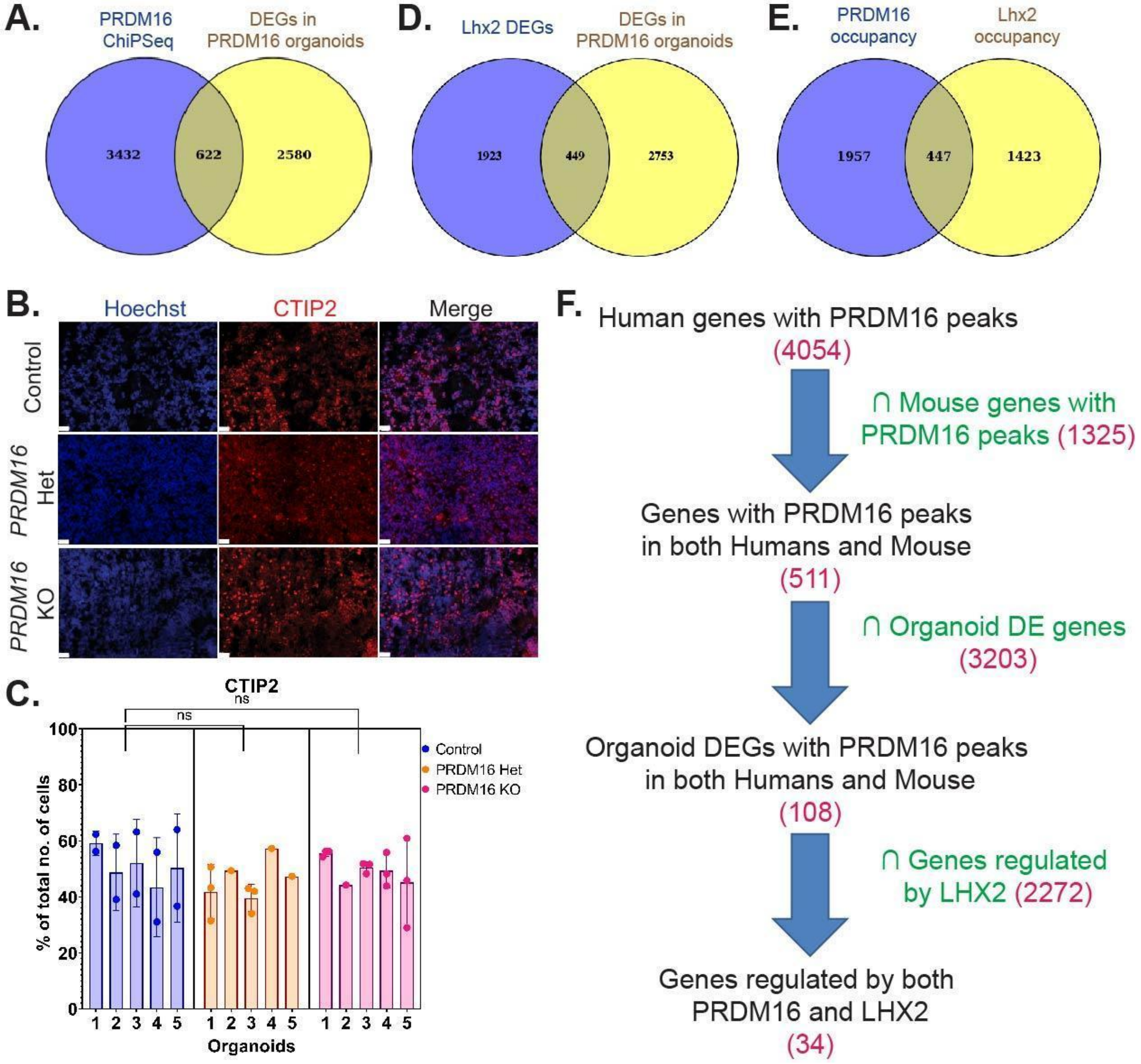


**Supplementary Fig. 4**

**A.** Venn diagram plot identifies genes that are occupied by PRDM16 in the ChIP-seq data and set of DEGs upon loss of PRDM16 in organoids **B.** Immunohistochemistry on 12 µm cryosections of day 32 control and PRDM16 mutant cortical organoids showing the cortical neuronal markers CTIP2 (imaged using 25X objective lens, scale bar represents 20 µm) **C.** Percentage of the total number of cells that are CTIP2$^+$ neurons, in day 32 control and PRDM16 mutant cortical organoids (Nested One-Way ANOVA, n = 5, α = 0.05, each data point on the graph represents an image of the organoid along its z-axis) **D.** Venn diagram plot identifies dysregulated genes that are common upon loss of *Lhx2,* and loss of *PRDM16* in the organoids **E.** Venn diagram plot identifies a common set genes that are occupied by LHX2 and PRDM16 in the mouse. **F.** Sequential filtration of genes identifies a common gene regulatory network possibly regulated by both LHX2 and PRDM16.

**Supplementary Fig. 5**

### A. Human *PRDM16*

| Rank | Motif | Name | P-value | log P-pvalue | q-value (Benjamini) | # Target Sequences with Motif | % of Targets Sequences with Motif |
|---|---|---|---|---|---|---|---|
| 1 | 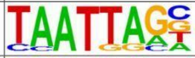 | Lhx2(Homeobox)/HFSC-Lhx2-ChIP-Seq(GSE48068)/Homer | 1e-501 | -1.154e+03 | 0.0000 | 2495.0 | 34.88% |
| 2 | 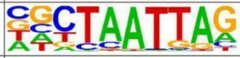 | LHX9(Homeobox)/Hct116-LHX9.V5-ChIP-Seq(GSE116822)/Homer | 1e-472 | -1.088e+03 | 0.0000 | 2852.0 | 39.87% |
| 3 | 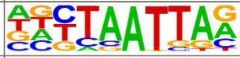 | Lhx1(Homeobox)/EmbryoCarcinoma-Lhx1-ChIP-Seq(GSE70957)/Homer | 1e-461 | -1.062e+03 | 0.0000 | 2570.0 | 35.92% |
| 4 | 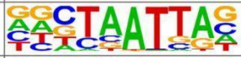 | En1(Homeobox)/SUM149-EN1-ChIP-Seq(GSE120957)/Homer | 1e-448 | -1.034e+03 | 0.0000 | 3327.0 | 46.51% |
| 5 | 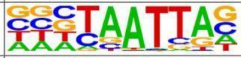 | DLX2(Homeobox)/BasalGanglia-Dlx2-ChIP-seq(GSE124936)/Homer | 1e-422 | -9.720e+02 | 0.0000 | 2965.0 | 41.45% |
| 6 | 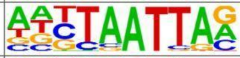 | Lhx3(Homeobox)/Neuron-Lhx3-ChIP-Seq(GSE31456)/Homer | 1e-400 | -9.217e+02 | 0.0000 | 3080.0 | 43.05% |
| 7 | 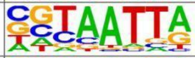 | DLX5(Homeobox)/BasalGanglia-Dlx5-ChIP-seq(GSE124936)/Homer | 1e-371 | -8.545e+02 | 0.0000 | 1890.0 | 26.42% |
| 8 | 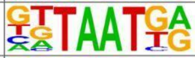 | Nkx6.1(Homeobox)/Islet-Nkx6.1-ChIP-Seq(GSE40975)/Homer | 1e-342 | -7.891e+02 | 0.0000 | 3922.0 | 54.82% |
| 9 | 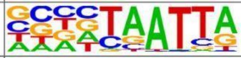 | DLX1(Homeobox)/BasalGanglia-Dlx1-ChIP-seq(GSE124936)/Homer | 1e-330 | -7.605e+02 | 0.0000 | 2583.0 | 36.11% |
| 10 | 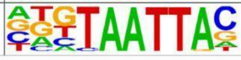 | Dlx3(Homeobox)/Kerainocytes-Dlx3-ChIP-Seq(GSE89884)/Homer | 1e-317 | -7.302e+02 | 0.0000 | 1620.0 | 22.64% |
| 141 | 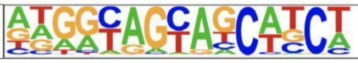 | PRDM9(Zf)/Testis-DMC1-ChIP-Seq (GSE35498)/Homer | 1e-3 | -9.060e+00 | 0.0004 | 370.0 | 5.17% |
| 173 | 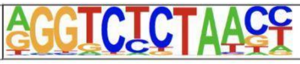 | PRDM14(Zf)/H1-PRDM14-ChIP-Seq (GSE22767)/Homer | 1e-2 | -5.182e+00 | 0.0143 | 210.0 | 2.94% |

### B. Mouse PRDM16

| Rank | Motif | Name | P-value | log P-pvalue | q-value (Benjamini) | # Target Sequences with Motif | % of Targets Sequences with Motif |
|---|---|---|---|---|---|---|---|
| 1 | 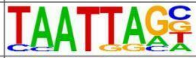 | Lhx2(Homeobox)/HFSC-Lhx2-ChIP-Seq(GSE48068)/Homer | 1e-229 | -5.287e+02 | 0.0000 | 876.0 | 40.17% |
| 2 | 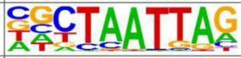 | LHX9(Homeobox)/Hct116-LHX9.V5-ChIP-Seq(GSE116822)/Homer | 1e-217 | -5.017e+02 | 0.0000 | 991.0 | 45.44% |
| 3 | 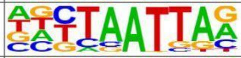 | Lhx1(Homeobox)/EmbryoCarcinoma-Lhx1-ChIP-Seq(GSE70957)/Homer | 1e-215 | -4.973e+02 | 0.0000 | 895.0 | 41.04% |
| 4 | 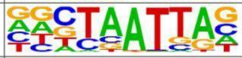 | En1(Homeobox)/SUM149-EN1-ChIP-Seq(GSE120957)/Homer | 1e-181 | -4.175e+02 | 0.0000 | 1082.0 | 49.61% |
| 5 | 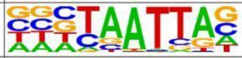 | DLX2(Homeobox)/BasalGanglia-Dlx2-ChIP-seq(GSE124936)/Homer | 1e-180 | -4.156e+02 | 0.0000 | 970.0 | 44.48% |
| 6 | 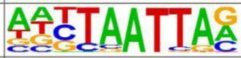 | Lhx3(Homeobox)/Neuron-Lhx3-ChIP-Seq(GSE31456)/Homer | 1e-170 | -3.917e+02 | 0.0000 | 1027.0 | 47.09% |
| 7 | 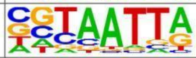 | DLX5(Homeobox)/BasalGanglia-Dlx5-ChIP-seq(GSE124936)/Homer | 1e-161 | -3.728e+02 | 0.0000 | 634.0 | 29.07% |
| 8 | 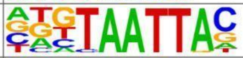 | Dlx3(Homeobox)/Kerainocytes-Dlx3-ChIP-Seq(GSE89884)/Homer | 1e-155 | -3.592e+02 | 0.0000 | 561.0 | 25.72% |
| 9 | 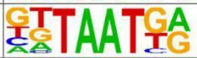 | Nkx6.1(Homeobox)/Islet-Nkx6.1-ChIP-Seq(GSE40975)/Homer | 1e-148 | -3.417e+02 | 0.0000 | 1264.0 | 57.96% |
| 10 | 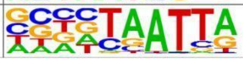 | DLX1(Homeobox)/BasalGanglia-Dlx1-ChIP-seq(GSE124936)/Homer | 1e-138 | -3.201e+02 | 0.0000 | 833.0 | 38.19% |

**A.** Top 10 motifs and the two PRDM family motifs identified in the human PRDM16 ChIP-seq data **B.** Top 10 motifs identified in the mouse PRDM16 ChIP-seq data on regions that are conserved across the mouse and human peaks.
